# Investigating the Impact of Hypoxia and Syncytialization on Lipid Nanoparticle-Mediated mRNA Delivery to Placental Cells

**DOI:** 10.1101/2025.06.23.661191

**Authors:** Rachel E. Young, Tara Vijayakumar, Darji Krisha, Samuel Hofbauer, Mohamad-Gabriel Alameh, Drew Weissman, Rachel Riley

## Abstract

Placental dysfunction leads to pregnancy-related disorders that affect up to 15% of pregnancies. Several of these, such as preeclampsia, are symptomatically managed but have no curative treatments other than preterm delivery. Placental dysfunction arises from improper placental development, leading to restricted blood vessel formation and a hypoxic placental microenvironment. The development of placental therapeutics is challenging due to the complex physiology that enables the placenta to have precise control over endocytosis and transport. Here, we use a simple culture system that combines hypoxia and trophoblast syncytialization to model the functional syncytiotrophoblast layer of the placenta under hypoxic stress. Using this model, we evaluate the impact of hypoxia on lipid nanoparticle (LNP)-mediated mRNA delivery. Our data shows that hypoxia hinders syncytiotrophoblast formation *in vitro*. Despite this, LNP delivery to syncytiotrophoblasts increases protein translation and secretion, particularly under hypoxic conditions. Further, we show delivery of a therapeutic mRNA, placental growth factor (PlGF), to syncytiotrophoblasts in hypoxia, which restored diminished PlGF levels back to normoxic controls. These findings provide an LNP platform for efficient mRNA delivery to hypoxic trophoblasts and demonstrate the importance of considering hypoxia towards the development of drug delivery platforms for placental therapeutics.

**Translational Impact Statement:** This study investigates lipid nanoparticle (LNP)-mediated mRNA delivery to placental cells cultured in a simple hypoxia model, simulating the microenvironment of the placenta during pregnancy complications such as preeclampsia. We provide insights essential for developing LNP drug delivery platforms for treating disorders of the placenta. These findings will guide future design of LNP-based treatments for delivery to the diseased, hypoxic placenta, minimizing systemic exposure and improving outcomes in maternal and fetal health for conditions lacking effective, targeted interventions.

## 1. Introduction

Throughout pregnancy, the placenta regulates immune activity, acts as a protective barrier for the fetus, and facilitates oxygen and nutrient exchange for fetal development. Improper formation and development of the placenta drives the onset and progression of placental dysfunction-related disorders, such as preeclampsia, Hemolysis, Elevated Liver enzymes, Low Platelet count (HELLP) syndrome, intrauterine and fetal growth restriction.^1–3^ Approximately 10 to 15% of pregnancies are affected by these conditions;^3^ but there are limited prophylactic or treatment options for severe cases. The only curative option for severe cases is to induce preterm delivery, which can have dire consequences on infant health when early in gestation (<28 weeks).^4–6^ Recently, nanoparticle-based strategies, including lipid nanoparticles (LNPs) have emerged to deliver therapeutic nucleic acids to the placenta to promote vascularization. Here, we sought to investigate interactions between LNPs and the placenta to inform the design and development of next-generation drug delivery platforms to address placental dysfunction-related disorders.

Abnormal placental development, including insufficient trophoblast invasion and uterine spiral artery remodeling, often coinciding with increased oxidative stress, is hallmark across various placental dysfunction-related disorders.^1^ Early in pregnancy (<10 weeks), cytotrophoblasts differentiate into extravillous trophoblasts and syncytiotrophoblasts. Extravillous trophoblasts are a highly invasive phenotype that remodel the uterine spiral arteries to establish blood flow to the placenta.^7^ Syncytiotrophoblasts are formed through the fusion of cytotrophoblasts to form multinucleated cells that act as a semipermeable barrier between the maternal and fetal blood circulation (Figure 1A).^7–10^ At this early stage of pregnancy, low oxygen tension and expression of hypoxia-inducible factor-1α (HIF-1α) are essential for embryo implantation and proper placental development;^11–13^ however, prolonged hypoxia and expression of HIF-1α in the 2^nd^ and 3^rd^ trimester stimulates the transcription of genes implicated in placental dysfunction-related disorders.^14–23^ Specifically, HIF-1α drives transcription of soluble fms-like tyrosine kinase-1 (sFlt-1),^24,25^ which sequesters placental growth factor (PlGF) in the placenta, limiting angiogenesis and vascular development.^17–21^

**Figure 1.**
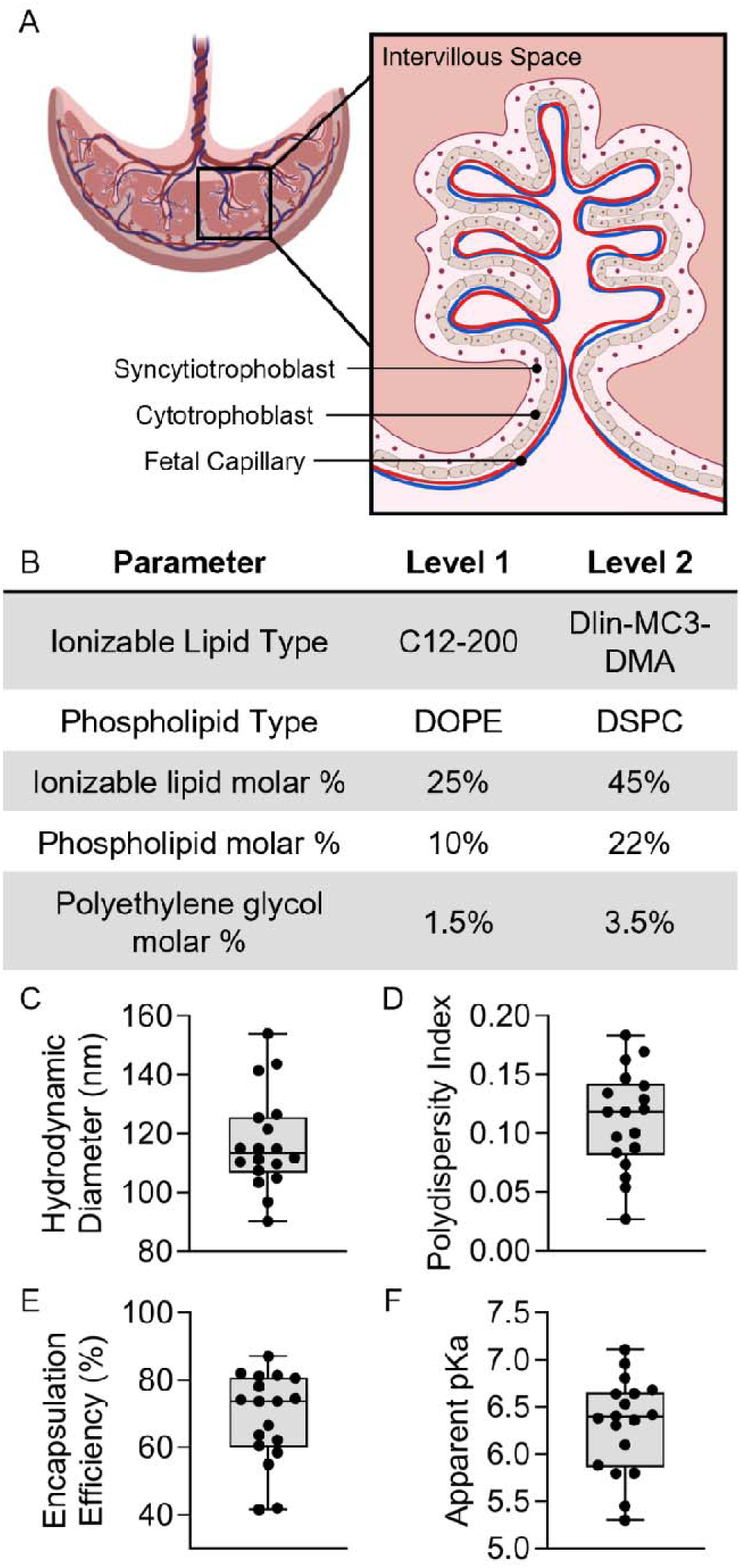
Placental structure and Luc-LNP library characterization. (A) Schematic of trophoblast layers in placental villi. Cytotrophoblasts fuse to form the syncytiotrophoblast, a multinucleated cell layer. (B) Definitive Screening Design parameters for the Luc-LNP library. (C) Hydrodynamic diameter, (D) polydispersity index, (E) encapsulation efficiency, and (F) apparent pKa of Luc-LNPs in the library. Each dot represents an LNP in the library.

In this work, we characterize the effects of hypoxia on trophoblasts in a straightforward *in vitro* model, and we use this model to assess the impact of hypoxia on LNP-mediated drug delivery. Although placental hypoxia plays a critical role in placental dysfunction, there are minimal studies evaluating how this diseased microenvironmental cue impacts placental-drug interactions. In other diseases, such as cancer, hypoxia has been shown to influence these interactions, leading to decreased nucleic acid delivery compared to normoxia.^26–29^ We and others have developed nanoparticle-based strategies, most commonly LNPs, to deliver therapeutic nucleic acids to the placenta to treat placental dysfunction-related disorders, such as preeclampsia.^34–45^ In prior work, we developed LNPs that deliver a range of nucleic acids to trophoblasts representative of the 3^rd^ trimester and to mouse placentas during early and late gestation.^34,35^

Here, we evaluate how LNP composition drives nucleic acid delivery to trophoblasts that represent different trimesters of pregnancy under normal and restrained oxygen environments. This provides the first study that evaluates LNP interactions with hypoxic syncytiotrophoblasts as the disease-relevant trophoblast subtype. Our approach combines a hypoxic cell culture model with forskolin-induced syncytialization to represent the hypoxic placenta in placental dysfunction-related disorders. We demonstrate that hypoxia significantly alters trophoblasts by increasing HIF-1α expression, decreasing PlGF secretion, and increasing cell growth. Further, hypoxia hinders forskolin-induced syncytialization, which in turn increases LNP-mediated mRNA delivery compared to normoxia. Lastly, we demonstrate the relevance of LNPs for protein replacement therapy in placental dysfunction-related disorders by delivering placental growth factor (PlGF) mRNA, which is a potential therapeutic molecule for preeclampsia. Together, the data presented here advances LNP design for nucleic acid delivery to the hypoxic placenta during different trimesters and demonstrates the importance of considering hypoxia to achieve drug delivery in the diseased placenta. Ultimately, this work lays the foundation for advancing LNPs for placental dysfunction-related disorders, which addresses a critical unmet need in maternal-fetal medicine.

## 2. Results

### 2.1 LNP Library Formulation and Characterization

LNPs are multilamellar nanoparticles comprised of four lipid components – ionizable lipids, phospholipids, cholesterol, and poly(ethylene) glycol (PEG) lipid conjugates – complexed with nucleic acids.^46,47^ Modulating the types and amounts of each lipid component within LNPs enables preferential delivery to specific tissues,^48–51^ an approach we used to achieve preferential delivery to mouse placentas in a prior study.^34^ Here, we use the same LNP library – created using a Design of experiments (DOE)^34^ – to examine how LNP design influences delivery to 1^st^ and 3^rd^ trimester trophoblasts under normoxia or hypoxia.

We encapsulated luciferase mRNA into LNPs (Luc-LNPs) as it is detectable and quantifiable using a plate reader. The Luc-LNP library consisted of 18 LNPs formulated with either C12-200 (C12) or DLin-MC3-DMA (MC3) as the ionizable lipid and 1,2-dioleoyl-sn-glycero-3-phosphoethanolamine (DOPE) or 1,2 distearoyl-sn-glycero-3-phosphocholine (DSPC) as the phospholipid (Figure 1B, Table S1). Additionally, we varied the molar percentages of ionizable lipid (25-45%), phospholipid (10-22%), and (1,2-dimyristoyl-sn-glycero-3-phosphoethanolamine-N-[methoxy(polyethylene glycol)-2000] (ammonium salt)) (DMPE-PEG) (1.5-3.5%) used to make the Luc-LNPs (Figure 1B, Table S1). The remaining molar percentage (to add up to 100%) in each Luc-LNP was cholesterol. The Luc-LNP library had hydrodynamic diameters ranging from 90.3 nm-153.9 nm (Figure 1C, Table S1) with low polydispersity below 0.2 for all formulations (Figure 1D, Table S1). The Luc-LNP library had mRNA encapsulation efficiencies ranging from 41.5%-87.1% relative to the amount of mRNA added during formulation (Figure 1E, Table S1). Apparent pKa was determined by measuring surface ionization with 6-(p-toluidinyl)naphthalene-2-sulfonic acid (TNS) assays, which determines the pH at which half of the ionizable lipids are protonated to induce endosomal escape and cytoplasmic mRNA delivery.^52,53^ The apparent pKa values ranged from 5.3-7.1 (Figure 1F, Table S1). Analyzing these LNP characteristics as responsive variables in our DOE, we found that the type of ionizable lipid was a significant factor affecting LNP apparent pKa (Table S2). We used this library to assess luciferase mRNA delivery to multiple trophoblast cell lines and under normoxia or hypoxia, as described below.

### 2.2 LNP Design Drives Delivery to Trophoblasts

To assess LNP delivery *in vitro*, we utilized three trophoblast cell lines: HTR8/SVneo (referred to as HTR8 herein), JAR, and the b30 subclone of BeWo cells (referred to as BeWo herein). HTR8 cells are representative of 1^st^ trimester trophoblasts,^54^ and JAR and BeWo cells are choriocarcinoma cells representative of 3^rd^ trimester placentas.^55–57^ We were interested in evaluating these three cell lines since they are most commonly used to represent early- and late-stage pregnancy, respectively.^24,58^ We treated cells with each Luc-LNP in the library at 0.75 nM mRNA or PBS for 24 hours. The top LNP formulation for each cell line varied, although similar formulations were high-performing across all cell lines. In HTR8 cells, LNP 8 had the highest increase in luminescence and LNPs 4, 5, 8, 10, and 14 all exhibited significantly increased fold change in luminescense compared to PBS-treated cells (Figure 2A, Table 1). In JAR cells, LNP 5 was the top-performing formulation and LNPs 4, 5, 8, and 10 were all significantly increased compared to PBS-treated cells (Figure 2B, Table 1). Lastly, BeWo cells treated with LNP 10 had the highest fold change in luminescence, and LNPs 5, 8, and 10 were significantly increased compared to PBS-treated cells (Figure 2C, Table 1). We examined metabolic activity as proxy for cell viability following delivery with the top LNPs from each cell line. LNPs 8 and 5 reduced metabolic activity by <10% in HTR8 and JAR cells at all doses (Figure S1A-B). In BeWo cells, LNP 10 reduced metabolic activity by 27% only at the highest dose, 0.75 nM (Figure S1C). Given the 95,535-fold increase in luciferase expression at this dose (Figure 2C), lower doses can be used for delivery to avoid any toxicity.

**Figure 2.**
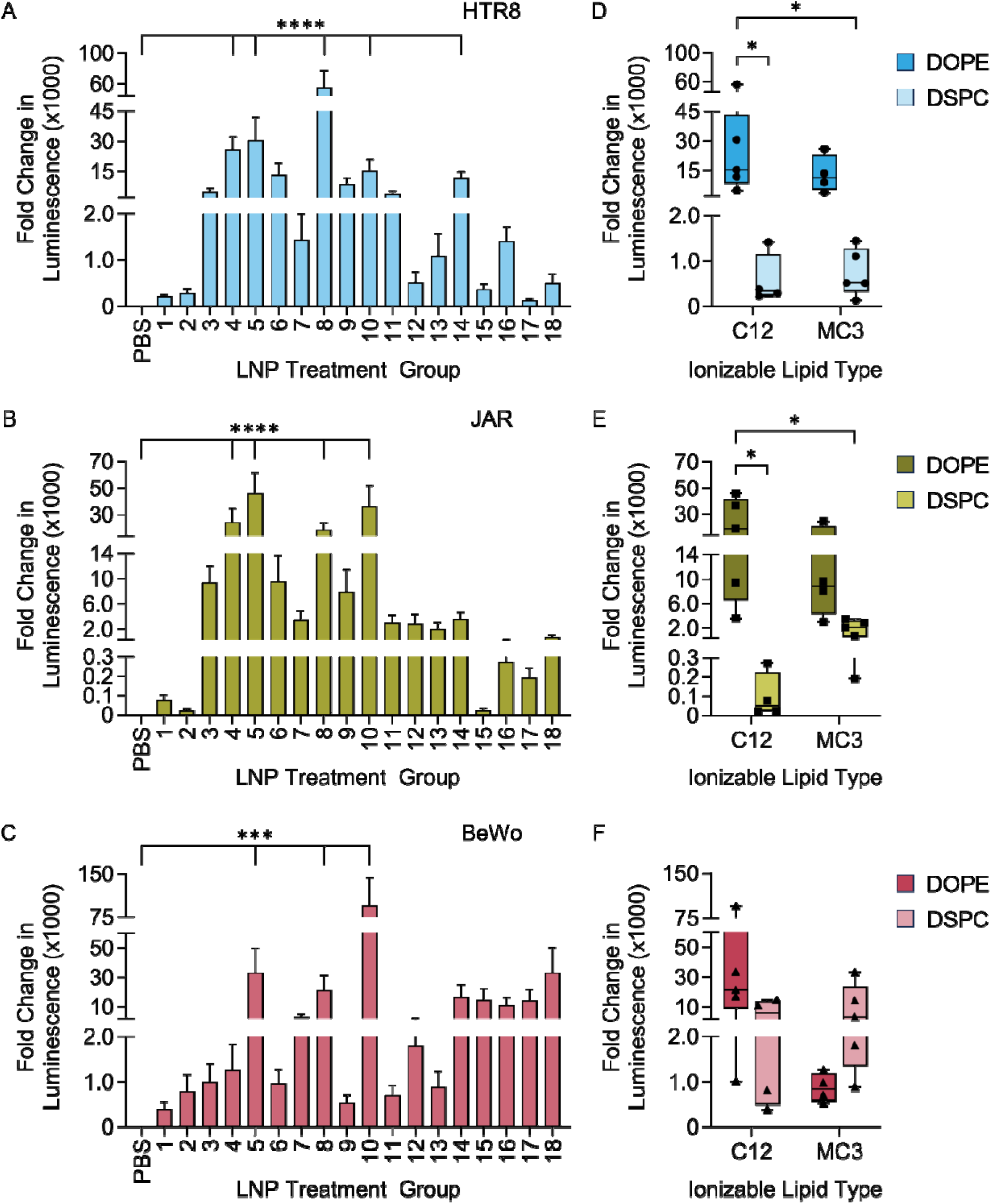
In vitro evaluation of the Luc-LNP library in trophoblast cell lines. Luminescence from (A) HTR8, (B) JAR, and (C) BeWo cells treated with Luc-LNPs was calculated as a fold change over the PBS-treated group within each cell line;*p<0.05, **p<0.01, ***p<0.001, ****p<0.0001 by Kruskal-Wallis test with Dunn’s post hoc test indicating signficance of each Luc-LNP compared to PBS. Luminescence from (D) HTR8, (E) JAR, and (F) BeWo cells grouped by type of ionizable lipid and phospholipid (right); *p<0.05 by two-way ANOVA with post hoc Tukey. Each marker represents an individual Luc-LNP from the library.

To determine the important design parameters for each cell line, we grouped luciferase data by lipid type (Figure 2D-F). LNPs containing DOPE had significantly increased luminescence fold change in HTR8 and JAR cells compared to LNPs containing DSPC, regardless of ionizable lipid type (Figure 2D-E). In the DOE analysis, the type of phospholipid was a signficant factor affecting LNP delivery in HTR8 (p=0.0012) and JAR (p=0.013) cells, with DOPE yielding the strongest luciferase expression. Furthermore, the amount of phospholipid in the formulation significantly affected LNP delivery in HTR8 cells (p=0.016), with a higher molar percentage of phospholipid yielding stronger luciferase expression. DOE analysis and two-way ANOVA tables are provided in Supplement Tables S3-S7. These results suggest that DOPE, as the phospholipid in Luc-LNPs, drives delivery in JAR and HTR8 cells. Comparatively, there was no statistical difference in fold change in luminescence based on lipid type in BeWo cells (Figure 2F).

Additionally, the DOE analysis did not identify any significant factors influencing LNP delivery to BeWo cells. However, the three LNPs with the highest statistical significance for delivery (LNP 5, 8, and 10) contain C12 and DOPE, suggesting that these two lipids are important for delivery to these cells. This agrees with our prior work demonstrating the importance of ionizable lipid and phospholipid type in BeWo cells.^34^

### 2.3 Hypoxic Culture Alters Trophoblast Behavior

Due to the biological complexity of oxygen tension throughout various stages of pregnancy, we first assessed how hypoxia alters trophoblast growth and hypoxia-related biomarker expression. Towards this goal, we compared trophoblasts cultured in a low oxygen environment (1% O_2_, referred to as hypoxia) to trophoblasts cultured in a room oxygen environment (21% O_2_, referred to as normoxia). To establish a hypoxic environment, we used a cell culture chamber purged with low oxygen gas comprised of 5% CO_2_, 1% O_2_, and balance nitrogen (Figure S2A). The chamber reached an equilibrium concentration of 1.39% O_2_ after 3 minutes (Figure S2B).

We measured HIF-1α expression in cells accumulating over time as an indicator of cellular response to hypoxia.^18,59^ As expected, hypoxia increased cumulative HIF-1α expression in all cell lines compared to normoxia over 72 hours (Figure 3A, Table S8-S9). We also assessed how hypoxia alters secretion of PlGF from cells, as reduced PlGF is characteristic of trophoblasts in hypoxia and in preeclamptic placentas.^60^ Culture in hypoxia decreased PlGF secretion from JAR and BeWo cells at all time points compared to normoxia (Figure 3B). Alternatively, PlGF secretion from HTR8 cells did not change between hypoxia and normoxia (Figure 3B). This was expected because PlGF levels begin to increase at the end of the 1^st^ trimester, peaking around 30 weeks;^61^ as a 1^st^ trimester cell line, HTR8 cells are not expected to produce high levels of PlGF. For all cell lines, the cumulative amount of PlGF in the culture media increased over time (Figure 3B). However, hypoxic culture decreased the rate of PlGF secretion (Figure 3B, Table S10-S11). In addition to changes to HIF-1α and PlGF, we also measured how hypoxia changes cell growth using MTS assays. We plotted and fitted a growth curve with a simple linear regression, which revealed that hypoxia significantly increases the growth rate of all three trophoblast cell lines (Figure 3C-E). Together, these results demonstrate that hypoxia alters trophoblast activity and supports our motivation to study how these physiological changes impact nanoparticle-mediated drug delivery in the hypoxic, diseased placenta.

**Figure 3.**
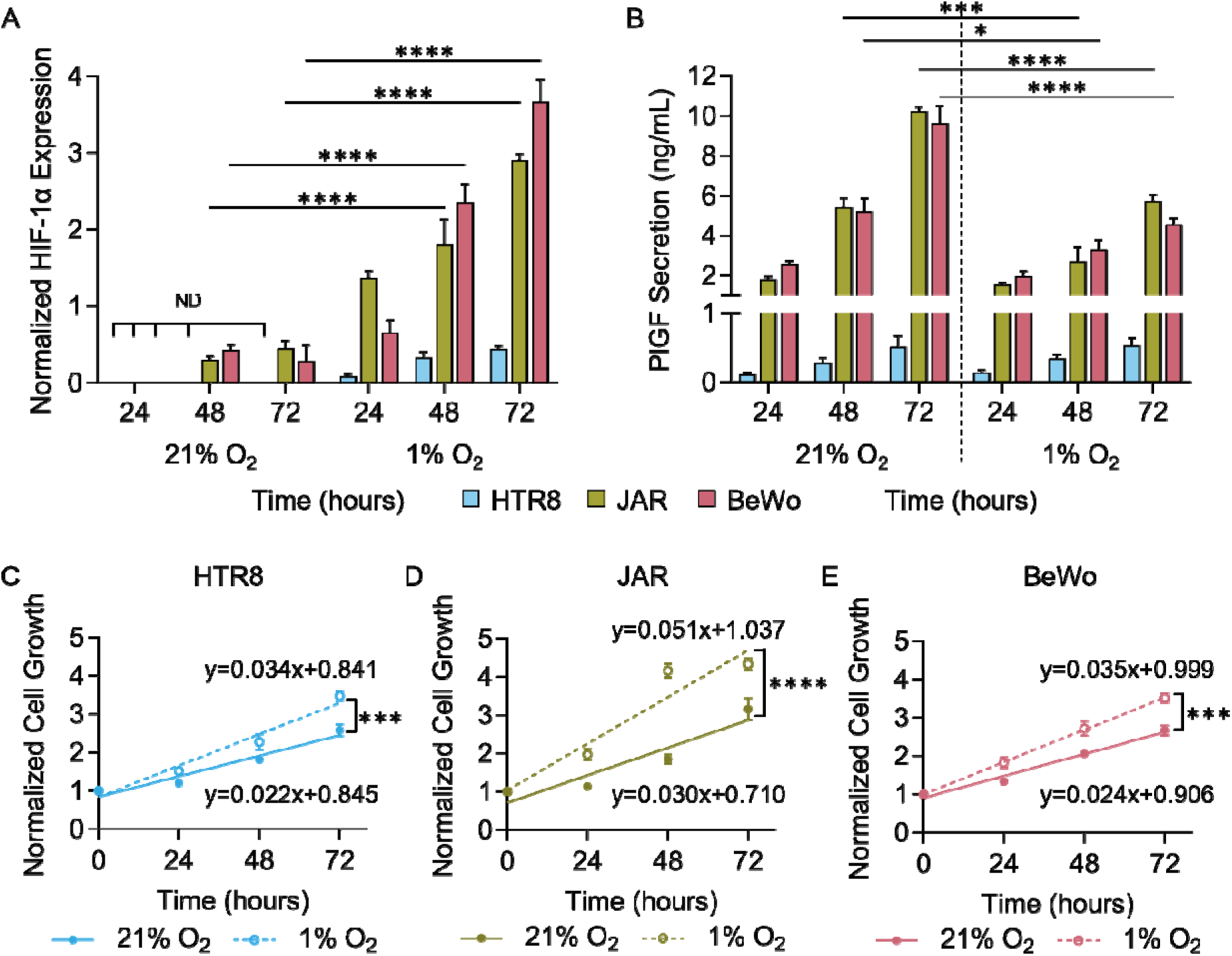
Cellular response to culture in under hypoxic (1% O2) conditions. (A) HIF-1α expression in cells and (B) PlGF secretion from cells cultured in normoxia (21% O2) or hypoxia (1% O2) at 24, 48, and 72 hours. “ND” is not detectable. *p<0.05, ***p<0.001 and ****p<0.0001 by two-way ANOVA with post hoc Tukey. (C) HTR8, (D) JAR, and (E) BeWo growth curves during cell culture in normoxia and hypoxia over time. ***p<0.001 and ****p<0.0001 by Analysis of Covariance (ANCOVA). Additional statistical analyses are provided in Tables S8-S11.

### 2.4 Hypoxia Increases LNP Delivery to Trophoblasts

Next, we assessed how hypoxia influences LNP-mediated mRNA delivery to trophoblasts using LNPs 5, 8, and 10 formulated with GFP mRNA (GFP-LNPs). These LNP designs were the top LNP candidates for each cell line in the library screen and they all contain C12 and DOPE as ionizable lipid and phospholipid, respectively (Figure 2, Table S1). In these experiments, GFP mean fluorescence intensity (MFI) was measured by flow cytometry in trophoblasts cultured in normoxia or hypoxia 2 and 24 hours following GFP-LNP administration. In all the cell lines, MFI was higher at 24 hours compared to 2 hours, which we expected because longer administration of LNPs allows for increased uptake and translation of mRNA (Figure 4). Further, the difference in MFI between cells cultured in hypoxia and normoxia at 24 hours is greater than the difference at 2 hours (Figure 4), indicating that the longer treatment time is more appropriate to evaluate the impact of hypoxia on LNP delivery.

**Figure 4.**
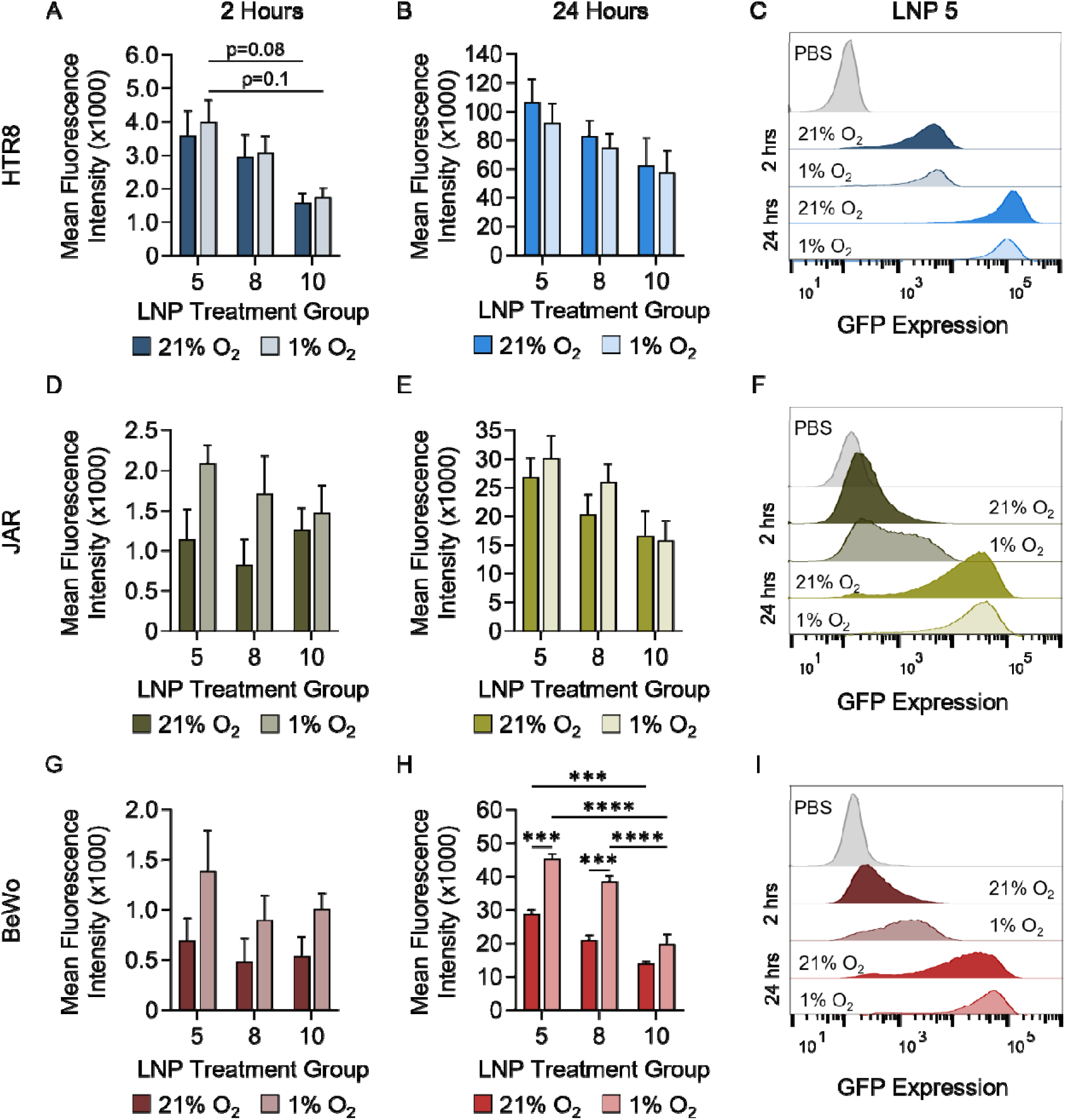
GFP-LNP delivery to cells cultured under normoxic and hypoxic conditions. GFP MFI following treatment with GFP-LNPs 5, 8, and 10 with autofluorescence subtracted from (A-C) HTR8, (D-F) JAR, and (G-I) BeWo cells cultured in normoxia (21% O_2_) or hypoxia (1% O_2_). Data was quantified by flow cytometry 2 hours (left) and 24 hours (center) after GFP-LNP treatment. Representative histogram data from LNP 5 are shown in the right column. ***p<0.001, ****p<0.0001 by two-way ANOVA with post hoc Tukey. Additional analyses of LNP design features are provided in Tables S12-S17.

Overall, GFP-LNP 5 yielded the highest MFI in all cell lines compared to GFP-LNPs 8 and 10 at both oxygen conditions and timepoints (Figure 4). Additionally, HTR8 cells overall had the highest delivery compared to the other cell lines (Figure 4). Importantly, culture in hypoxia did not significantly affect GFP-LNP delivery in HTR8 and JAR cells based on GFP MFI (Figure 4A-F). Alternatively, hypoxic culture significantly impacted delivery to BeWo cells. For example, at 24 hours following treatment with GFP-LNPs 5 and 8, BeWo cells cultured in hypoxia had significantly increased MFI compared to normoxia (Figure 4H). Additional statistical analyses comparing LNP design factors are presented in Supplemental Tables S11-S16. We also examined BeWo cell metabolic activity following delivery with GFP-LNPs, which revealed no significant difference between cells in normoxia and hypoxia (Figure S3). These results suggest that hypoxia increases uptake of LNPs and translation of mRNA in BeWo cells, and this difference in delivery is not due to alterations to cellular metabolism following LNP delivery.

### 2.5 Hypoxia Inhibits Trophoblast Syncytialization

The experiments described above suggest that placental hypoxia, which is characteristic in placental dysfunction-related disorders, likely alters the interactions of LNPs with trophoblasts. This demonstrates the importance of studying delivery under diseased microenvironmental cues. To create a more physiologically-relevant culture system to study LNP interactions with the placenta, we next aimed to understand how hypoxia impacts syncytialization. Syncytialization refers to the fusion of trophoblasts that form the functional outer barrier of the placenta (Figure 1A). The syncytiotrophoblast layer controls and facilitates exchange and transport of endogenous compounds or drug molecules, and drives placental metabolism.^10,62,63^ Further, syncytiotrophoblasts are major producers of hormones, such as human chorionic gonadotropin (hCG), and angiogenic factors to support and maintain pregnancy.^62,63^ Thus, the design of nanoparticle-based delivery platforms, including LNPs, needs to consider delivery of the therapeutic cargo to these specialized, functional cells. For this reason, we next assessed the impact of hypoxia on syncytialized trophoblasts.

We cultured HTR8, JAR, and BeWo cells in hypoxia or normoxia and chemically induced trophoblast syncytialization with forskolin.^64–67^ Forskolin is a cyclic AMP (cAMP) analog, commonly used to induce trophoblast fusion *in vitro*, creating a syncytiotrophoblast-like phenotype.^58,68^ Following forskolin treatment, we measured tight junction formation via immunofluorescence staining of zonula occludens-1 (ZO-1) expression in the cells (Figure 5A-D). Here, a decrease in ZO-1 expression following forskolin treatment indicates cellular fusion during syncytialization.^58,69^ HTR8 cells did not have ZO-1 expression in control (DMSO) or forskolin-treated groups in normoxia or hypoxia (Figure 5A-B). HTR8 cells are an invasive, 1^st^ trimester extravillous trophoblast (EVT) cell line;^54,70–72^ therefore, we did not expect tight junctions to form or cellular fusion to occur after forskolin treatment. In comparison, forskolin decreased ZO-1 expression in JAR cells cultured in hypoxia moreso than in normoxia (Figure 5A,C). This suggests that hypoxia sensitizes JAR cells to forskolin-induced cell fusion. In BeWo cells, forskolin treatment significantly reduced ZO-1 expression in both normoxia and hypoxia compared to cells not treated with forskolin (Figure 5D). This suggests that hypoxia does not significantly alter forskolin-induced cellular fusion of BeWos, prompting us to explore additional markers of syncytialization to fully assess the impact of hypoxia on these cells.

**Figure 5.**
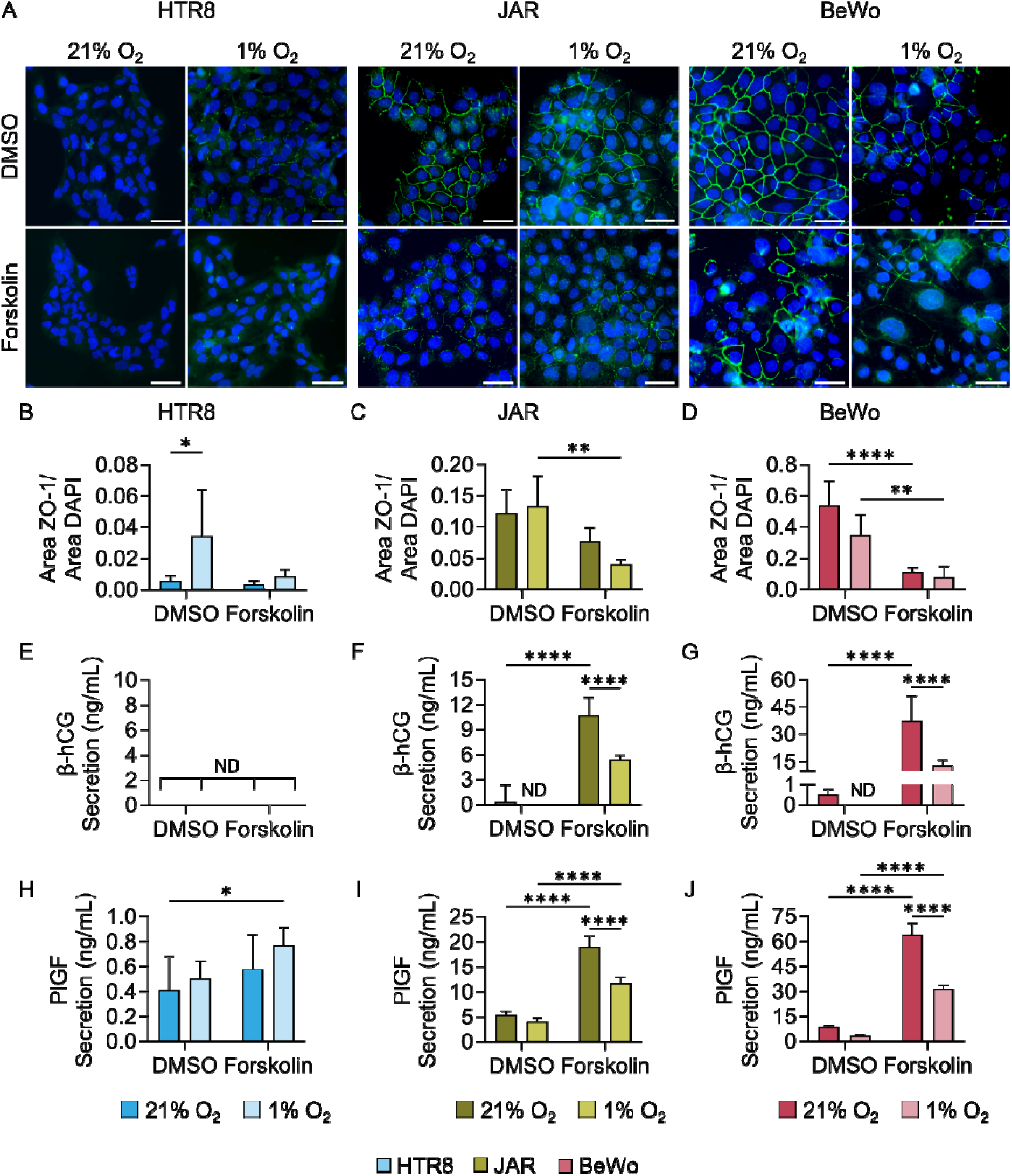
Impact of hypoxia on trophoblast syncytialization. (A) ZO-1 (green) and nuclei (blue) expression in HTR8 (left), JAR (center), and BeWo (right) cells treated with DMSO (control) or forskolin and cultured in normoxia (21% O_2_) or hypoxia (1% O_2_). Scale bar=50 µm. (B-D) Quantification of ZO-1 area normalized to nuclei area in each image (n=5). (E-G) β-human chorionic gonadotropin (β-hCG) and (H-J) placental growth factor (PlGF) secretion from each cell line. *p<0.05, **p<0.01, ***p<0.001, and ****p<0.0001 by two-way ANOVA with post hoc Tukey. Additional analysis shown in Tables S18-S21.

To further assess the impact of hypoxia on syncytialization, we also measured β-hCG and PlGF secretion from trophoblasts (Figure 5E-J). The secretion of β-hCG from HTR8 cells was undetectable by the assay kit, likely because EVTs are not the main trophoblast subtype secreting β-hCG during pregnancy (Figure 5E).^73^ However, hypoxia significantly increased PlGF secretion from HTR8 cells compared to normoxia (Figure 5H), although the magnitude of its secretion was lower than JAR or BeWo cells. In both JAR and BeWo cells cultured in hypoxia and normoxia, forskolin increased β-hCG and PlGF secretion (Figure 5F, G, I, J). However, in the forskolin-treated cells, β-hCG and PlGF secretion was signficantly decreased in hypoxia compared to normoxia (Figure 5F, G, I, J). We ran a 2-way ANOVA on this data, which identified that forskolin treatment and hypoxic culture together drive hCG and PlGF secretion from JAR and BeWo cells (Table S18-S21). This suggests that hypoxia inhibits the ability of forskolin to induce secretion of β-hCG and PlGF, both of which are markers of syncytialization. This demonstrates the importance of considering hypoxia in the placenta when developing therapeutics, as cellular behavior is greatly impacted by chronic hypoxia.

### 2.6 Syncytialization Alters LNP Interactions with Trophoblasts

As described above, our data suggests that BeWos undergo increased syncytialization compared to the other two cell lines in normoxia, as represented by high β-hCG and PlGF secretion. For this reason, we utilized BeWo cells to investigate how syncytialization alters LNP interactions. For these experiments, we used GFP-LNP 5 based on its ability to deliver GFP mRNA to these cells (Figure 4). BeWos were cultured in hypoxia or normoxia for 48 hours prior to treatment with DMSO or forskolin for another 48 hours. Next, cells were treated with GFP-LNP 5 for 24 hours prior to analysis by flow cytometry. The syncytialized (forskolin-treated) BeWos cultured in hypoxia had significantly increased MFI following LNP delivery compared to all other experimental groups (Figure 6A). This indicates that syncytialization increases LNP delivery, in agreement with prior literature.^41^ However, this data also indicates that this increased delivery is amplified when the syncytialized cells are cultured under hypoxia. This finding contrasts prior studies, which found that hypoxia decreases nanoparticle uptake in cancer cells.^26^ The potential for increased delivery to trophoblasts in hypoxia underscores the importance of considering this microenvironmental cue during preclinical drug development, as increased uptake in the diseased placenta, compared to the healthy placenta, could lead to undesired toxicities to the placenta and/or fetus.

**Figure 6.**
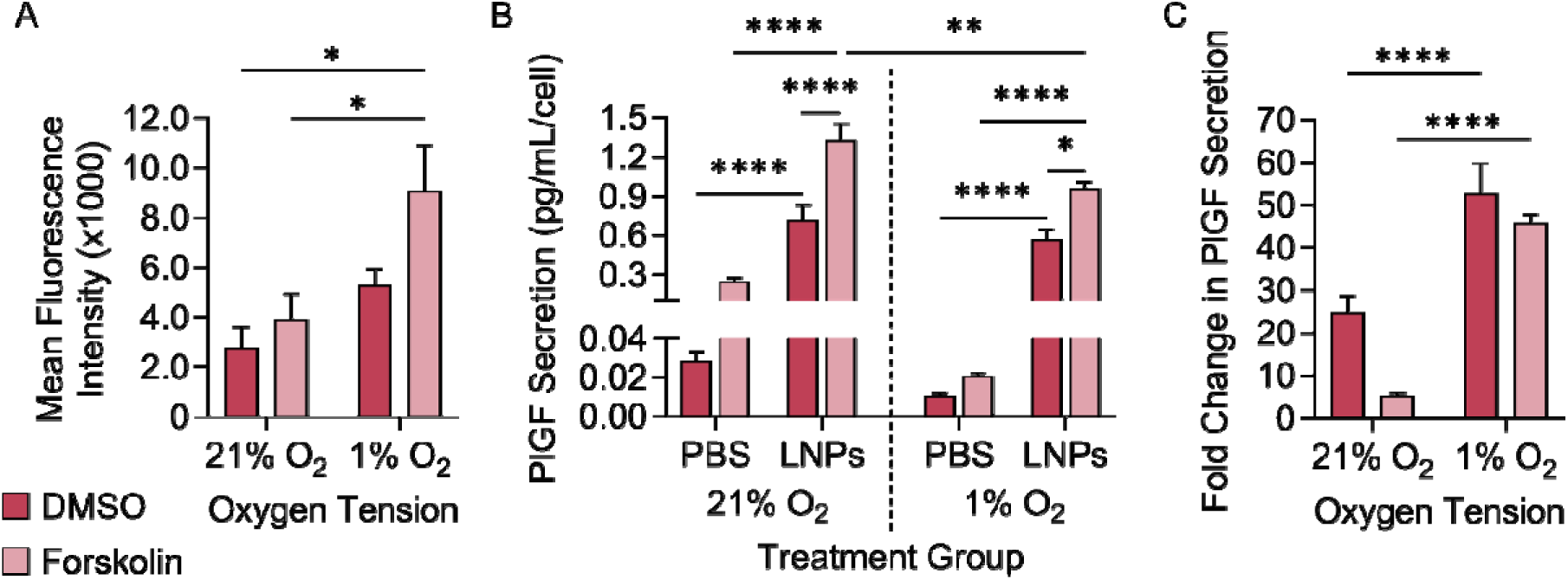
LNP delivery to syncytialized and non-syncytialized BeWo cells cultured under hypoxic and normoxic conditions. (A) GFP MFI from LNP-treated syncytialized or non-syncytialized BeWo cells under each oxygen condition. In these experiments, BeWo cells were pre-treated with DMSO or forskolin to induce syncytialization and cultured under normoxia (21% O_2_) or hypoxia (1% O_2_) prior to GFP-LNP delivery, and GFP MFI was quantified by flow cytometry after 24 hours. Data shown is raw MFI with autofluorescence subtracted. *p<0.05 by two-way ANOVA with post hoc Tukey. (B) Normalized PlGF secretion 24 hours after treatment with PlGF-LNPs. In this data, raw PlGF secretion was normalized by the number of cells at the time of analysis to account for cell growth 24 hours after LNP treatment. *p<0.05, **p<0.01, ***p<0.001, ****p<0.0001 by three-way ANOVA with post hoc Tukey. (C) Fold-change of PlGF secretion from LNP-treated cells relative to PBS-treated cells 24 hours after treatment with PlGF-LNPs. ****p<0.0001 by two-way ANOVA with post hoc Tukey. Additional statistical analyses shown in Tables S22-S24.

To demonstrate that our findings hold relevance towards a protein replacement therapy, we replaced the GFP mRNA with placental growth factor (PlGF) mRNA (PlGF-LNPs). PlGF is normally secreted by trophoblasts and plays a critical role in supporting angiogenesis and vascularization in the placenta. Further, PlGF levels in the placenta are decreased in placental dysfunction-related disorders compared to healthy pregnancies – demonstrating its potential as a target for protein replacement therapy.^74–77^ We treated cells with PlGF-LNPs and compared the resultant PlGF secretion to PBS-treated controls. Since our data shows that hypoxia increases cellular growth (Figure 3E), we normalized PlGF secretion in these experiments to the number of cells at the time of sample collection (Figure 6B). All cells treated with PlGF-LNPs had increased PlGF secretion compared to PBS-treated groups cultured under the same oxygen or syncytialization conditions (Figure 6B). However, culture under each condition revealed trends regarding how oxygen or syncytialization impacts PlGF-LNP delivery. First, our data shows that syncytialized cells had increased PlGF secretion compared to non-syncytialized cells in both normoxia and hypoxia (Figure 6B). This indicates that the increased LNP uptake in syncytialized cells corresponds to better secretion of therapeutic proteins, supporting the use of syncytialized cells in preclinical drug development.

Second, hypoxia decreased PlGF secretion from syncytialized cells treated with PlGF-LNPs compared to normoxia (Figure 6B). Our prior data (Figure 3B) showed that hypoxia decreases endogenous PlGF secretion. Based on this, we aimed to determine if hypoxia decreases LNP-mediated PlGF secretion, or if this data reflects endogenous changes to secretion. We compared the fold change in PlGF secretion from LNP-treated and PBS-treated cells in the same oxygen and syncytialization conditions (Figure 6C). This allowed us to compare the increase in PlGF secretion from LNP-treated groups while accounting for basal levels. The fold change in PlGF secretion from cells in hypoxia was significantly higher than in normoxia (Figure 6C). This suggests that PlGF-LNPs increased PlGF secretion in cells cultured in hypoxia moreso than normoxia. Further, this difference is greater in syncytialized cells compared to non-syncytialized cells (Figure 6C). Together, this data indicates that syncytialization and hypoxia in combination increase PlGF-LNP delivery and subsequent PlGF secretion. Further, it shows that LNPs can restore PlGF levels secreted by syncytiotrophoblasts under hypoxic stress, setting the stage for future work delivering PlGF mRNA as a therapeutic in diseases implicated by placental hypoxia.

## 3. Discussion

The main goal of this study was to investigate how hypoxia, which is a common microenvironmental feature of placental dysfunction-related disorders, impacts nanoparticle-based drug delivery to the placenta. The insights reported herein provide a foundational understanding of how hypoxia alters trophoblast behavior, and how these changes in turn impact the efficiency of LNP-mediated mRNA delivery. More broadly, this work demonstrates the need to incorporate disease-relevant microenvironmental cues, such as hypoxia, into the preclinical development of drug delivery platforms.

As mentioned previously, the role of hypoxia changes as pregnancy progresses; hypoxia is required in early pregnancy, but a lack of reoxygenation in the 2^nd^ trimester yields undesired oxidative stress.^14–16^ Due to the variable role of hypoxia at different stages of pregnancy, we evaluated 1^st^ (HTR8) and 3^rd^ (JAR and BeWo) trimester trophoblast cell lines cultured in a 1% oxygen environment. Our data demonstrates that hypoxia increases cell growth and HIF-1α expression in all trophoblast cell lines. Increased expression of HIF-1α in the 2^nd^ and 3^rd^ trimesters is associated with placental dysfunction-related disorders.^78^ This increased HIF-1α expression is negatively correlated with PlGF levels in placental tissue and blood serum of preeclamptic patients,^79^ suggesting that PlGF secretion is dependent on oxidative stress. Further, experimentally-induced overexpression of HIF-1α in pregnant mice causes symptoms of preeclampsia, HELLP, and IUGR, such as increased blood pressure, proteinuria, liver enzymes, and resultant glomerular endotheliosis and thrombocytopenia.^21^ Our results with JAR and BeWo cells demonstrate decreased PlGF secretion in hypoxia, indicating that hypoxia causes these cells, which are representative of the 3^rd^ trimester placenta, to exhibit a disease-like phenotype consistent with clinical data.

To enhance the relevance of our hypoxic trophoblast culture system, we induced cellular fusion using forskolin as a model of syncytialization. The syncytiotrophoblast layer of the placenta is the main functional unit that, among other critical roles, controls the transport of molecules into the placenta to support fetal development.^62,63^ Insufficient syncytiotrophoblast formation contributes to placental dysfunction-related disorders and can hinder the proper transfer of nutrients through the placenta.^80^ When cultured in hypoxia, BeWos and cytotrophoblasts isolated from the human placenta show suppressed syncytialization,^81–83^ which is regulated via HIF signaling pathways.^59,83–85^ Additionally, syncytiotrophoblasts decrease endogenous secretion of PlGF in hypoxic culture.^60^ Our results corroborate these findings by showing that hypoxia reduces hCG and PlGF secretion in forskolin-treated cells compared to normoxia. This indicates that a hypoxic microenvironment in the 3^rd^ trimester placenta may hinder the formation of the syncytiotrophoblast, limiting PlGF secretion. This finding is critical both for understanding the pathophysiology of placental dysfunction-related disorders and towards the development of drug delivery platforms, as altered syncytiotrophoblast formation could significantly impact nanoparticle delivery efficiency to the placenta.

The data presented herein is the first to investigate how hypoxia, a key feature of placental-dysfunction related disorders, impacts LNP uptake and mRNA translation in trophoblasts. The impact of hypoxia on LNP uptake has been evaluated in other disease applications.^26,27^ For example, LNP-mediated mRNA delivery to glioblastoma cells cultured in hypoxia was decreased compared to normoxia, likely due to decreased cellular ATP concentrations.^26^ To the contrary, our data shows that hypoxia increases cellular growth and LNP-mediated mRNA delivery to trophoblasts. We hypothesize that this increased LNP delivery is a result of biological changes to endocytic pathways and cellular permeability, both of which are regulated by differential oxygen conditions.^86^ For example, prolonged hypoxia was found to increase endocytosis via caveolin-1 (CAV-1)-mediated pathways in trophoblasts, which may explain the increased LNP delivery in hypoxia seen herein.^87^ In addition to altering endocytic pathways, hypoxia and forskolin-induced syncytialization could increase cellular permeability, thereby impacting LNP delivery.^41,67,88–90^ This hypothesis is supported by our data showing decreased ZO-1 levels in syncytialized cells and those cultured under hypoxia, which correlates increased LNP-mediated mRNA delivery. Taken together, these data underscore the importance of utilizing disease-relevant microenvironmental conditions to evaluate the performance of drug delivery platforms, such as LNPs, for treating placental dysfunction-related disorders.

To extend our study toward investigating a potential therapeutic target for protein replacement therapy, we evaluated LNP-mediated delivery of PlGF mRNA to hypoxic syncytiotrophoblasts. PlGF drives angiogenesis in the placenta, is important for proper maintenance throughout pregnancy, and its expression is decreased in the preeclamptic placenta.^91–94^ Previous reports have demonstrated delivery of recombinant PlGF as a potential protein replacement therapy to decrease arterial blood pressure and restore angiogenic factor balance in animal models of preeclampsia, demonstrating its clinical relevance.^74–77^ Previously, we utilized LNPs to deliver PlGF mRNA to pregnant mice, inducing production of PlGF in the placental tissue and secretion into the maternal serum.^34^ Herein, we aimed to understand how LNP-mediated PlGF secretion is impacted in the hypoxic placenta using our hypoxic culture model. Culture in hypoxia decreased endogenous PlGF secretion in BeWo cells, but LNPs were able to restore PlGF back to the same levels seen in normoxia. This confirms that LNPs exhibit increased delivery to hypoxic syncytiotrophoblasts, and they hold potential for protein replacement therapy to support vascularization in the hypoxic, preeclamptic placenta.

Herein, we evaluated how hypoxia impacts LNP delivery to trophoblasts across various stages of pregnancy. We accomplished this by using an *in vitro* model that more closely replicates the hypoxic microenvironment in the diseased placenta compared to normal culture conditions. Together, our results demonstrate that hypoxia, particularly in combination with syncytialization, significantly alters the LNP interactions with trophoblasts. A limitation of this study is that it does not fully replicate the complex cellular processes that occur within the placenta, such as continuous blood flow and interactions between various cell types. Here, we used forskolin to induce syncytialization in trophoblast cell lines, which very common in the literature;^58,68^ however, expression of genes in trophoblast cell lines can differ from spontaneously fusing cytotrophoblasts *in vivo.*^95^ Additionally, the hypoxic culture model used here is more acute (2 days prior to treatment) than chronic hypoxia over several months seen in the preeclamptic placenta. Moving forward, we aim to expand our model to include primary trophoblasts isolated from human placentas and multiple layers of cells to more closely represent the complex placental architecture. However, the trophoblast culture system we developed herein revealed the importance of considering hypoxia in the development of drug delivery platforms. This culture system could be applied in the design and optimization of drug delivery platforms for placental dysfunction-related diseases and for future mechanistic investigation of the molecular pathways driving LNP uptake in hypoxic and syncytialized cells. Understanding how diseased microenvironmental cues, such as hypoxia, impact delivery will ultimately guide the rational design of advanced delivery platforms to improve maternal-fetal outcomes in complicated pregnancies.

## 4. Conclusions

In this study we examined LNPs for mRNA delivery to 1^st^ and 3^rd^ trimester trophoblasts cultured in hypoxia, which is a critical microenvironmental factor in placental dysfunction-related disorders. We demonstrated increased HIF-1α expression and decreased PlGF secretion from 3^rd^ trimester trophoblasts cultured in hypoxia, indicative of a disease-relevant microenvironment. Additionally, hypoxia decreased syncytiotrophoblast formation. LNP uptake was also increased in hypoxic and forskolin-treated BeWo cells, highlighting the importance of considering the cellular microenvironment when designing LNPs for placental therapies. Additionally, we showed that LNPs can successfully deliver PlGF mRNA to hypoxic trophoblasts, establishing a foundation for therapeutic mRNA strategies to restore placental function. Our simple trophoblast culture system could be used to examine the cellular and molecular signaling pathways during placental dysfunction-related disorders. These findings advance the understanding of LNP interactions with trophoblasts and provide a framework for optimizing mRNA delivery approaches for placental dysfunction-related disorders. Ultimately, this work contributes to the growing field of placental nanomedicine, addressing a critical unmet need in maternal-fetal medicine.

## 5. Materials and Methods

### 5.1 LNP Formulation

Ionizable lipids, including C12-200 and DLin-MC3-DMA (MC3), were purchased from MedChem Express. Other LNP components including cholesterol, 1,2 distearoyl-sn-glycero-3-phosphocholine (DPSC), 1,2-dioleoyl-sn-glycero-3-phosphoethanolamine (DOPE), and (1,2-dimyristoyl-sn-glycero-3-phosphoethanolamine-N-[methoxy(polyethylene glycol)-2000] (ammonium salt)) (DMPE-PEG) were purchased from Avanti Polar Lipids, Inc. Codon-optimized mRNA was prepared by in vitro transcription through a collaboration with the Engineered mRNA and Targeted Nanomedicine core facility at the University of Pennsylvania. Firefly luciferase, eGFP mRNA, and PlGF mRNA (transcript variant 1, NM_002632.6) were co-synthesized with 1-methylpseudouridine modifications, and co-transcriptionally capped using the CleanCap system (TriLink) and purified using cellulose based chromatography. Each LNP in the library was formulated via mixing with micropipettes, combining one volume of the lipid ingredients in ethanol to three volumes of mRNA in citrate buffer (pH 3). The lipid mixture for each LNP formulation contained various molar ratios of ionizable lipid:phospholipid:cholesterol:PEG, as indicated in Table S1. mRNA was diluted in citrate buffer to an mRNA:ionizable lipid ratio of 1:10 for each of the LNP formulations. Following mixing of the two phases, LNPs were dialyzed against PBS (pH 7.4) for 2 hours, sterile filtered using a 0.2 µm filters, and stored at 4°C until use.

### 5.2 LNP Characterization

Dynamic light scattering (DLS) measurements and mRNA encapsulation efficiency, as described below, were measured in triplicate for each LNP in the library. For DLS, each LNP was diluted 1:100 in deionized water in cuvettes and intensity measurements were run on a Malvern Zetasizer Nano ZS (Malvern Panalytical). The encapsulation efficiency of each LNP formulation was calculated using QuantiFluor® RNA System (Promega) as previously described.^36^ Briefly, LNPs were diluted 1:100 in 1X TE buffer in two microcentrifuge tubes per LNP formulation. 1% v/v Triton X-100 (Thermo Scientific) was added to one of the tubes and both were heated to 37°C and shaken at 600 RPM for 5 minutes, followed by cooling to room temperature for 10 minutes. LNP samples and RNA standards were plated in triplicate in black 96-well plates and the fluorescent reagent was added per manufacturer instructions. Fluorescent intensity was read on the plate reader (excitation, 492 nm; emission, 540 nm). Background signal was subtracted from each well and triplicate wells for each LNP were averaged. RNA content was quantified by comparing samples to the standard curve, and encapsulation efficiency (%) was calculated according to the equation 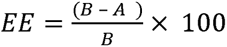, where A is the RNA content in samples without Triton X-100 treatment (intact LNPs) and B is the RNA content in samples treated with Triton X-100 (lysed LNPs).

The apparent pKa of LNPs was quantified via [6-(p-toluidinyl)naphthalene-2-sulfonic acid] (TNS, Thomas Scientific) assays, as previously described.^96^ Briefly, a buffer solution of 150 mM sodium chloride, 20 mM sodium phosphate, 20 mM ammonium acetate, and 25 mM ammonium citrate (VWR Chemicals BDH) was prepared and separated into pH-adjusted tubes ranging from pH 2 to pH 12 in increments of 0.5 pH. 2.5 µL of each LNP formulation was combined with 125 µL of each pH-adjusted solution in black 96-well plates in triplicate. TNS was added to each well for a final TNS concentration of 6 µM and the fluorescence intensity was read on a plate reader (Molecular Devices) (excitation, 322 nm; emission, 431 nm). Fluorescence versus pH was plotted, and apparent pKa was calculated as the pH corresponding to 50% of its maximum value, representing 50% protonation.

### 5.3 Normoxic or Hypoxic Cell Culture

Three trophoblast cell lines, HTR8/SVneo (termed “HTR8” herein), JAR, and the b30 subclone^68^ of the BeWo choriocarcinoma cell line (termed “BeWos” herein) were used to investigate mRNA delivery with the LNP library. HTR8 and JAR cells were cultured in RPMI 1640 with 2.05 mM L-glutamine supplemented with 10% fetal bovine serum (Avantor) and 1% penicillin/streptomycin (VWR). BeWo cells were cultured in F-12K Nutrient Mixture (Kaighn’s Mod.) with L-glutamine (Corning Inc.) supplemented with 10% fetal bovine serum and 1% penicillin/streptomycin. Prior to experimentation, all cells were cultured under normal oxygen cultures (termed “normoxia” herein) in a room air incubator set at 37°C supplemented with 5% CO_2_. For experiments using low oxygen cultures (termed “hypoxia” herein), a hypoxic cell culture chamber (StemCell Technologies) was purged with low oxygen gas (1% O2, 5% CO2, and 94% N2) for 5 minutes. Oxygen concentration was confirmed using a Go Direct^TM^ oxygen sensor (Vernier Software & Technology). The cell culture chamber was then placed in a warm room with temperature set at 37°C.

### 5.4 LNP Delivery to Trophoblasts

HTR8, JAR, and BeWo cells were plated at 20,000 cells per well in 96-well plates with 200 µL of complete media in triplicate for each Luc-LNP formulation. After 4 hours, cells were treated with Luc-LNPs diluted in sterile PBS at 100 ng mRNA/well (0.75 nM) or sterile PBS as the negative control. Luciferase expression was analyzed after 24 hours per manufacturer instructions (Promega). Cells were washed with sterile PBS and 20 µL of 1x lysis buffer was added to each well. After 10 minutes of incubation at room temperature, cells were centrifuged at 12,000 xg for 2 minutes, and lysates were plated into white 96-well plates. 100 µL of luciferase assay substrate was added to each well and the luminescent signal was quantified using the plate reader. The average luminescent signal from each group was normalized to untreated cells and reported as the fold change in luminescence. Statistical analysis of luciferase expression from the Luc-LNP library screen was conducted (see “Statistical Analysis” section below) to determine significance.

To assess metabolic activity as an indicator of cell viability, HTR8, JAR, and BeWo cells were plated as described above and treated with 25 – 100 ng mRNA/well (0.1875 nM – 0.75 nM) of each top Luc-LNP formulation. After 24 hours, cells were assayed using the CellTiter 96® AQueous One Solution Cell Proliferation Assay (MTS; Promega) according to manufacturer instructions. Briefly, after the Luc-LNP treatment, 20 µL of the MTS reagent was added to each well. Cells were incubated for 2 hours at 37°C and absorbance at 490 nm was read on the plate reader. The average absorbance of wells containing no cells was subtracted as background from each well. The absorbance signal from each group was normalized to untreated cells and reported as the fold change in absorbance.

### 5.5 Impact of Hypoxia on Production of HIF-1α, PlGF and Cell Growth

To assess the impact of hypoxia on trophoblasts in culture, we quantified HIF-1α inside cells and PlGF secreted from cells. To assess intracellular HIF-1α, HTR8, JAR, and BeWo cells were plated at 150,000 cells per well in 6-well plates with 2 mL of complete media and cultured under normoxic or hypoxic conditions (see above) for a total of 72 hours. At 24, 48, and 72 hours after plating, different wells for each timepoint were washed with PBS and treated with 100 µL of 1x lysis buffer supplemented with 1x protease and phosphatase inhibitor cocktail (Pierce Biotechnology) to release protein content. To assess secreted PlGF, cells were plated at 50,000 cells per well in 24-well plates with 1 mL of complete media and cultured under normoxic or hypoxic conditions for 72 hours. At 24, 48, and 72 hours after plating, cell culture media was collected from different wells for each timepoint. Lysates and cell culture media were centrifuged at 10,000 xg for 10 minutes to remove cell debris and the supernatant was stored at −80°C until analysis. Lysates and cell culture media were assayed for HIF-1α and PlGF, respectively, using an enzyme-linked immunosorbent assay (ELISA) per manufacturer instructions (HIF-1α, ThermoFisher; PlGF, Rockland Immunochemicals, Inc.). Sample absorbance at 450 nm were read on the plate reader and were compared to a standard curve to calculate HIF-1α concentration inside cells and PlGF in the culture media.

To assess the impact of hypoxia on cell growth, HTR8, JAR, and BeWo cells were plated at 5,000 cells per well in 96-well plates with 200 µL of complete media and cultured under normoxic or hypoxic conditions for a total of 72 hours. At 24, 48, and 72 hours after plating, using different wells for each timepoint, MTS reagent was added to each well. Cells were incubated for 2 hours at 37°C in their respective oxygen condition and absorbance at 490 nm was read on the plate reader. The average absorbance of wells containing no cells at each time point was subtracted as background from each well. The absorbance signal from each group was normalized to the initial absorbance for each cell line under each oxygen condition. Data was reported as the normalized absorbance, plotted against time, and fit with a simple linear regression (see “Statistical Analysis”).

### 5.6 LNP Delivery to Hypoxic Cells

HTR8, JAR, and BeWo cells were plated at 150,000 cells per well in 6-well plates with 2 mL of complete media and cultured under normoxic or hypoxic conditioned (see above). After 48 hours of culture, cell culture media was replaced, and cells were treated with each top GFP-LNP formulation, dosed at 300 ng mRNA per well (4.5 nM). During LNP treatment, cells were kept under normoxic or hypoxic culture. After 2 or 24 hours, cells were detached from the culture plate with 0.25% trypsin with 0.2 g/L EDTA (Cytiva, Logan, UT) for ~2 minutes. Then, cells were diluted with complete media and collected into tubes for centrifugation. Cells were washed 3 times in Fluorescence Activated Cell Sorting (FACS) buffer (Rockland Immunochemicals) and analyzed based on GFP mean fluorescence intensity (MFI) with autofluorescence subtracted using a SA3800 Spectral Analyzer (Sony). Data was analyzed using FlowJo 10.10.0 software (BD Biosciences).

To assess metabolic activity as an indicator of cell viability, HTR8, JAR, and BeWo cells were 5,000 cells per well in 96-well plates with 200 µL of complete media and cultured under normoxic or hypoxic conditioned (see above). Cells were treated with 25 ng mRNA/well (4.5 nM) of each top GFP-LNP formulation. After 24 hours, cells were assayed using the CellTiter 96® AQueous One Solution Cell Proliferation Assay (MTS; Promega) according to manufacturer instructions. Briefly, after GFP-LNP treatment, 20 µL of the MTS reagent was added to each well. Cells were incubated for 2 hours at 37°C and absorbance at 490 nm was read on the plate reader. The average absorbance of wells containing no cells was subtracted as background from each well. The absorbance signal from each group was normalized to untreated cells and reported as the fold change in absorbance.

### 5.7 Trophoblast Syncytialization and Fluorescence Imaging

The impact of syncytialization on cell phenotype was assessed by protein production and fluorescence imaging. First, HTR8, JAR, and BeWo cells were plated at 50,000 cells per well in 24-well plates in 1 mL of complete media. After 4 hours, the culture media was replaced with complete media supplemented with 50 µM of forskolin (Enzo Life Sciences) diluted in dimethyl sulfoxide (DMSO, VWR), or volume-matched DMSO, and cells were cultured under normoxic or hypoxic conditions. The final concentration of DMSO in media was 0.5% (v/v). After 48 hours, culture media was collected and centrifuged at 10,000 xg for 10 minutes to remove debris. Beta human chorionic gonadotropin (β-hCG) and PlGF in the culture media was measured by ELISA per manufacturer instructions (β-hCG, DRG International Inc.; PlGF, Rockland Immunochemicals, Inc.).

For fluorescence imaging, HTR8, JAR, and BeWo cells were plated at 10,000 cells per well in 8-well Nunc® Lab-Tek™ chambered coverglasses (Thermo Scientific) in 200 µL of complete media. Each experimental group was plated in duplicate. Cells were allowed to adhere to the slide in normoxia for 4 hours. Next, the culture media was replaced with complete media supplemented with 50 µM of forskolin diluted in DMSO or volume-matched DMSO, and cultured under normoxic or hypoxic conditions. The final concentration of DMSO in media was 0.5% (v/v). After 48 hours of culture, cells were washed with PBS and fixed to the coverglasses with 3.7% formaldehyde (VWR) for 20 minutes at room temperature. Cells were washed 3 times with PBS and permeabilized with 0.25% Tween-20 (VWR) for 5 minutes at room temperature. Cells were washed with PBS and blocked with 3% bovine serum albumin (BSA, VWR) for 1 hour at room temperature. Cells were stained with ZO-1 Monoclonal Antibody (ZO1-1A12), Alexa Fluor™ 488 (339188, 0.5 mg/mL, Invitrogen) diluted 1:100 in 3% BSA for 2.5 hours at room temperature. Cells were washed 3 times with PBS and nuclei were stained with bisBenzimide H 33258 trihydrochloride (Hoechst 33258, 10 mg/mL in water, Biotium) diluted 1:1000 in PBS for 10 minutes at room temperature. Cells were washed 5 times in PBS and stored in PBS protected from light at 4 °C prior to imaging.

All imaging was conducted with a Nikon eclipse Ti2 inverted microscope with a Plan Apo λ 20x/0.75 objective and a Nikon DS-Qi2 camera with 3 s and 250 ms exposure for FITC and DAPI filters, respectively. ZO-1 and nuclei were imaged with the FITC and DAPI filter cubes, respectively. Five ROI’s were imaged per well to a total of 10 images for quantification. The area of ZO-1 and nuclei fluorescence within each ROI were quantified using ImageJ (National Institutes of Health). The area of ZO-1 fluorescense in the ROI was divided by the area of nuclei fluorescense in the same ROI for each image to account for the number of cells in each image. The mean and standard error of the mean of the area ratio was reported for each experimental group.

### 5.8 LNP Delivery to Syncytiotrophoblasts in Hypoxia

For GFP mRNA delivery, BeWo cells were plated at 50,000 cells per well in a 6-well plate in 2 mL of complete media. For PlGF mRNA delivery, BeWo cells were plated at 12,500 cells per well in a 24-well plate in 1 mL of complete media. In both experiments, cells were cultured under normoxic or hypoxic conditions for 48 hours. Next, cells were syncytialized following the protocol described above – culture media was replaced with complete media supplemented with 50 µM of forskolin or volume-matched DMSO – and cells were kept under normoxic or hypoxic culture conditions. After 48 hours, culture media was replaced with fresh forskolin- or DMSO-supplemented media and LNPs were added. Cells were treated with LNPs dosed at 200 ng GFP mRNA per well (3.0 nM) or 100 ng PlGF mRNA per well (3.0 nM) for 24 hours. After 24 hours, cells in the GFP mRNA delivery experiment were prepared and analyzed by flow cytometry, as described above. In the PlGF mRNA delivery experiment, the cell culture media was collected, and cells were counted. The cell culture supernatant was assayed for PlGF expression using an ELISA, as described above.

### 5.9 Statistical Analysis

The fold change in luminescense for each cell line following delivery with the Luc-LNP library is represented as the mean with standard error of the mean (SEM) with n=3 biological replicates. The data was assessed by Design of Experiments (DOE) analysis and Kruskal-Wallis tests. DOE analysis was conducted in JMP Pro 18 (SAS Institute, Inc.) software using the fit definitive screening platform. JMP Pro 18 uses effective model selection for DSDs to identify design variables as active main or pairwise interactions when the p-value computed using the t Ratio and degrees of freedom for error is less than 0.05. After active effects are identified in the Combined Model Parameter Estimates report, a standard least squares fit is applied to obtain the significant effects in the fit model. Additionally, the data was assessed for normality using D’Agostino-Pearson omnibus (K2), Anderson-Darling (A2*), Shapiro-Wilk (W), and/or Kolmogorov-Smirnov (distance) in Prism 10.4.0 software (GraphPad). The data was non-normal; therefore, a Kruskal-Wallis test followed by Dunn’s multiple comparisons test for each Luc-LNP in the library was performed. The fold change in luminescense data grouped by lipid type was assessed by Ordinary Two-Way ANOVA followed by Tukey’s method for pairwise comparisons of the means. Statistical significance for all Luc-LNP library analysis was determined at p<0.05 (*), p<0.01 (**), p<0.001 (***), or p<0.0001 (****).

Prism 10.4.0 software was used to perform all other statistical analysis. All results are represented in the figures as the mean with standard error of the mean (SEM) unless otherwise indicated. All experiments have n=3 biological replicates unless otherwise indicated. An Ordinary Two-Way ANOVA followed by Tukey’s method for pairwise comparisons of the means was used to analyze HIF-1α expression and PlGF secretion from cells cultured in hypoxia and normoxia over time. A simple linear regression was used to fit cell growth curves in hypoxia and normoxia over time and an Analysis of Covariance (ANCOVA) was used to determine statistical differences in the slope of the regression lines. GFP-LNP delivery to cells cultured in hypoxia and normoxia was assessed by an Ordinary Two-Way ANOVA followed by Tukey’s method for pairwise comparisons of the means. This statistical test was used to investigate the effect of LNP formulation and oxygen condition on MFI at 2 and 24 hours following GFP-LNP delivery. ZO-1 expression, β-hCG and PlGF secretion, and GFP-LNP delivery to cells treated with forskolin or DMSO and cultured in hypoxia or normoxia were assessed by an Ordinary Two-Way ANOVA followed by Tukey’s method for pairwise comparisons of the means. This statistical test was used for ZO-1 expression to investigate the effect of forskolin treatment and oxygen condition on the cells. PlGF secretion from cells following PlGF-LNP delivery to cells was assessed by an Ordinary Three-Way ANOVA followed by Tukey’s method for pairwise comparisons of the means. The fold change in PlGF secretion from cells was calculated by dividing the PlGF secretion from PlGF-LNP-treated groups by PlGF secretion from PBS-treated groups in the same oxygen or syncytialization conditions. The fold change was assessed by Ordinary Two-Way ANOVA followed by Tukey’s method for pairwise comparisons of the means. Statistical significance for all tests was determined at p<0.05 (*), p<0.01 (**), p<0.001 (***), or p<0.0001 (****).

## Supplementary Information

Supplemental images and tables are found in the attached supplementary information document.

## Acknowledgements

This work was supported by the Peter Joseph Pappas Fund through the Preeclampsia Foundation, the New Jersey Health Foundation (PC 44-22), the New Jersey Department of Health (COCR22PRG012), the National Science Foundation (NSF) Engineering Research Initiation Program (Award #2301919), and the NSF Graduate Research Fellowship Program (Award #2018266781).

## Literature Cited

1. Burton GJ, Redman CW, Roberts JM, Moffett A. Pre-eclampsia: pathophysiology and clinical implications. BMJ. Jul 15 2019;366:l2381. doi:10.1136/bmj.l2381

2. Barton JR, Sibai BM. Diagnosis and management of hemolysis, elevated liver enzymes, and low platelets syndrome. Clin Perinatol. Dec 2004;31(4):807–33, vii. doi:10.1016/j.clp.2004.06.008

3. Wardinger JE, Ambati S. Placental Insufficiency. StatPearls [Internet]. StatPearls Publishing LLC; 2022.

4. Odigboegwu O, Pan LJ, Chatterjee P. Use of Antihypertensive Drugs During Preeclampsia. Front Cardiovasc Med. 2018;5:50. doi:10.3389/fcvm.2018.00050

5. Berzan E, Doyle R, Brown CM. Treatment of preeclampsia: current approach and future perspectives. Curr Hypertens Rep. Sep 2014;16(9):473. doi:10.1007/s11906-014-0473-5

6. Armaly Z, Jadaon JE, Jabbour A, Abassi ZA. Preeclampsia: Novel Mechanisms and Potential Therapeutic Approaches. Front Physiol. 2018;9:973. doi:10.3389/fphys.2018.00973

7. Park JY, Mani S, Clair G, et al. A microphysiological model of human trophoblast invasion during implantation. Nature Communications. 2022/03/15 2022;13(1):1252. doi:10.1038/s41467-022-28663-4

8. Masserdotti A, Gasik M, Grillari-Voglauer R, et al. Unveiling the human fetal-maternal interface during the first trimester: biophysical knowledge and gaps. Front Cell Dev Biol. 2024;12:1411582. doi:10.3389/fcell.2024.1411582

9. Burton GJ, Fowden AL. The placenta: a multifaceted, transient organ. Philos Trans R Soc Lond B Biol Sci. Mar 5 2015;370(1663):20140066. doi:10.1098/rstb.2014.0066

10. Renaud SJ, Jeyarajah MJ. How trophoblasts fuse: an in-depth look into placental syncytiotrophoblast formation. Cell Mol Life Sci. Jul 20 2022;79(8):433. doi:10.1007/s00018-022-04475-z

11. Wakeland AK, Soncin F, Moretto-Zita M, et al. Hypoxia Directs Human Extravillous Trophoblast Differentiation in a Hypoxia-Inducible Factor-Dependent Manner. Am J Pathol. Apr 2017;187(4):767–780. doi:10.1016/j.ajpath.2016.11.018

12. Rosario GX, Konno T, Soares MJ. Maternal hypoxia activates endovascular trophoblast cell invasion. Dev Biol. Feb 15 2008;314(2):362–75. doi:10.1016/j.ydbio.2007.12.007

13. Zhao H, Wong RJ, Stevenson DK. The Impact of Hypoxia in Early Pregnancy on Placental Cells. Int J Mol Sci. Sep 7 2021;22(18) doi:10.3390/ijms22189675

14. Burton GJ, Cindrova-Davies T, Yung HW, Jauniaux E. HYPOXIA AND REPRODUCTIVE HEALTH: Oxygen and development of the human placenta. Reproduction. 2021;161(1):F53–F65. doi:10.1530/rep

15. Tong W, Giussani DA. Preeclampsia link to gestational hypoxia. J Dev Orig Health Dis. Jun 2019;10(3):322–333. doi:10.1017/S204017441900014X

16. Tianthong W, Phupong V. Serum hypoxia-inducible factor-1alpha and uterine artery Doppler ultrasound during the first trimester for prediction of preeclampsia. Sci Rep. Mar 23 2021;11(1):6674. doi:10.1038/s41598-021-86073-w

17. Pintye D, Sziva RE, Mastyugin M, et al. Nitroxide-HMP-Protects Human Trophoblast HTR-8/SVneo Cells from H(2)O(2)-Induced Oxidative Stress by Reducing the HIF1A Signaling Pathway. Antioxidants (Basel). Aug 8 2023;12(8)doi:10.3390/antiox12081578

18. Nevo O, Soleymanlou N, Wu Y, et al. Increased expression of sFlt-1 in in vivo and in vitro models of human placental hypoxia is mediated by HIF-1. Am J Physiol Regul Integr Comp Physiol. Oct 2006;291(4):R1085–93. doi:10.1152/ajpregu.00794.2005

19. Soleymanlou N, Jurisica I, Nevo O, et al. Molecular evidence of placental hypoxia in preeclampsia. J Clin Endocrinol Metab. Jul 2005;90(7):4299–308. doi:10.1210/jc.2005-0078

20. Tal R. The role of hypoxia and hypoxia-inducible factor-1alpha in preeclampsia pathogenesis. Biol Reprod. Jun 2012;87(6):134. doi:10.1095/biolreprod.112.102723

21. Tal R, Shaish A, Barshack I, et al. Effects of hypoxia-inducible factor-1alpha overexpression in pregnant mice: possible implications for preeclampsia and intrauterine growth restriction. Am J Pathol. Dec 2010;177(6):2950–62. doi:10.2353/ajpath.2010.090800

22. Burton GJ, Jauniaux E, Watson AL. Maternal arterial connections to the placental intervillous space during the first trimester of human pregnancy: The Boyd Collection revisited. Am J Obstet Gynecol. 1999;181(3):718–724. doi:10.1016/S0002-9378(99)70518-1

23. Jauniaux E, Watson AL, Hempstock J, Bao YP, Skepper JN, Burton GJ. Onset of maternal arterial blood flow and placental oxidative stress. A possible factor in human early pregnancy failure. American Journal of Pathology. 2000;157(6):2111–2122. doi:10.1016/S0002-9440(10)64849-3

24. Zhao J, Chow RP, McLeese RH, Hookham MB, Lyons TJ, Yu JY. Modelling preeclampsia: a comparative analysis of the common human trophoblast cell lines. FASEB BioAdvances. 2020;3(1):23–35. doi:10.1096/fba.2020-00057

25. Sasagawa T, Nagamatsu T, Yanagisawa M, Fujii T, Shibuya M. Hypoxia-inducible factor-1beta is essential for upregulation of the hypoxia-induced FLT1 gene in placental trophoblasts. Mol Hum Reprod. Nov 27 2021;27(12) doi:10.1093/molehr/gaab065

26. Ma Y, Fenton OS. An Efficacy and Mechanism Driven Study on the Impact of Hypoxia on Lipid Nanoparticle Mediated mRNA Delivery. J Am Chem Soc. May 24 2023;145(20):11375–11386. doi:10.1021/jacs.3c02584

27. Tiwade PB, Ma Y, VanKeulen-Miller R, Fenton OS. A Lung-Expressing mRNA Delivery Platform with Tunable Activity in Hypoxic Environments. J Am Chem Soc. Jun 26 2024;146(25):17365–17376. doi:10.1021/jacs.4c04565

28. Liu HM, Zhang YF, Xie YD, et al. Hypoxia-responsive ionizable liposome delivery siRNA for glioma therapy. Int J Nanomedicine. 2017;12:1065–1083. doi:10.2147/IJN.S125286

29. Li Y, Lu A, Long M, Cui L, Chen Z, Zhu L. Nitroimidazole derivative incorporated liposomes for hypoxia-triggered drug delivery and enhanced therapeutic efficacy in patient-derived tumor xenografts. Acta Biomater. Jan 1 2019;83:334–348. doi:10.1016/j.actbio.2018.10.029

30. Adams D, Gonzalez-Duarte A, O’Riordan WD, et al. Patisiran, an RNAi Therapeutic, for Hereditary Transthyretin Amyloidosis. N Engl J Med. Jul 5 2018;379(1):11–21. doi:10.1056/NEJMoa1716153

31. Riley RS, June CH, Langer R, Mitchell MJ. Delivery technologies for cancer immunotherapy. Nat Rev Drug Discov. Mar 2019;18(3):175–196. doi:10.1038/s41573-018-0006-z

32. Baden LR, El Sahly HM, Essink B, et al. Efficacy and Safety of the mRNA-1273 SARS-CoV-2 Vaccine. N Engl J Med. Feb 4 2021;384(5):403–416. doi:10.1056/NEJMoa2035389

33. Polack FP, Thomas SJ, Kitchin N, et al. Safety and Efficacy of the BNT162b2 mRNA Covid-19 Vaccine. N Engl J Med. Dec 31 2020;383(27):2603–2615. doi:10.1056/NEJMoa2034577

34. Young RE, Nelson KM, Hofbauer SI, et al. Systematic development of ionizable lipid nanoparticles for placental mRNA delivery using a design of experiments approach. Bioact Mater. Apr 2024;34:125–137. doi:10.1016/j.bioactmat.2023.11.014

35. Hofbauer S, Fink LA, Young RE, et al. Cytokine mRNA Delivery and Local Immunomodulation in the Placenta using Lipid Nanoparticles. bioRxiv. 2025; doi:10.1101/2025.02.07.637086

36. Riley RS, Kashyap MV, Billingsley MM, et al. Ionizable lipid nanoparticles for in utero mRNA delivery. Science Advances. 2021;7:1–15. doi:10.1126/sciadv.aba1028

37. Safford HC, Swingle KL, Geisler HC, et al. Orthogonal Design of Experiments for Engineering of Lipid Nan0oparticles for mRNA Delivery to the Placenta. Small. Aug 3 2023:e2303568. doi:10.1002/smll.202303568

38. Swingle KL, Hamilton AG, Safford HC, et al. Placenta-tropic VEGF mRNA lipid nanoparticles ameliorate murine pre-eclampsia. Nature. Dec 11 2024; doi:10.1038/s41586-024-08291-2

39. Swingle KL, Safford HC, Geisler HC, et al. Ionizable Lipid Nanoparticles for In Vivo mRNA Delivery to the Placenta during Pregnancy. J Am Chem Soc. Mar 1 2023;145(8):4691–4706. doi:10.1021/jacs.2c12893

40. Chaudhary N, Newby AN, Arral ML, et al. Lipid nanoparticle structure and delivery route during pregnancy dictate mRNA potency, immunogenicity, and maternal and fetal outcomes. Proc Natl Acad Sci U S A. Mar 12 2024;121(11):e2307810121. doi:10.1073/pnas.2307810121

41. Abostait A, Abdelkarim M, Bao Z, et al. Optimizing lipid nanoparticles for fetal gene delivery in vitro, ex vivo, and aided with machine learning. J Control Release. Oct 27 2024;376:678–700. doi:10.1016/j.jconrel.2024.10.047

42. Zhang B, Liang R, Zheng M, Cai L, Fan X. Surface-Functionalized Nanoparticles as Efficient Tools in Targeted Therapy of Pregnancy Complications. Int J Mol Sci. Jul 25 2019;20(15) doi:10.3390/ijms20153642

43. Zhang B, Tan L, Yu Y, et al. Placenta-specific drug delivery by trophoblast-targeted nanoparticles in mice. Theranostics. 2018;8(10):2765–2781. doi:10.7150/thno.22904

44. Li L, Li H, Xue J, Chen P, Zhou Q, Zhang C. Nanoparticle-Mediated Simultaneous Downregulation of Placental Nrf2 and sFlt1 Improves Maternal and Fetal Outcomes in a Preeclampsia Mouse Model. ACS Biomater Sci Eng. Oct 12 2020;6(10):5866–5873. doi:10.1021/acsbiomaterials.0c00826

45. Li L, Yang H, Chen P, et al. Trophoblast-Targeted Nanomedicine Modulates Placental sFLT1 for Preeclampsia Treatment. Front Bioeng Biotechnol. 2020;8:64. doi:10.3389/fbioe.2020.00064

46. Guimaraes PPG, Zhang R, Spektor R, et al. Ionizable lipid nanoparticles encapsulating barcoded mRNA for accelerated in vivo delivery screening. J Control Release. Dec 28 2019;316:404–417. doi:10.1016/j.jconrel.2019.10.028

47. Kulkarni JA, Darjuan MM, Mercer JE, et al. On the Formation and Morphology of Lipid Nanoparticles Containing Ionizable Cationic Lipids and siRNA. ACS Nano. May 22 2018;12(5):4787–4795. doi:10.1021/acsnano.8b01516

48. Dahlman JE, Kauffman KJ, Xing Y, et al. Barcoded nanoparticles for high throughput in vivo discovery of targeted therapeutics. Proc Natl Acad Sci U S A. Feb 21 2017;114(8):2060–2065. doi:10.1073/pnas.1620874114

49. Kauffman KJ, Webber MJ, Anderson DG. Materials for non-viral intracellular delivery of messenger RNA therapeutics. J Control Release. Oct 28 2016;240:227–234. doi:10.1016/j.jconrel.2015.12.032

50. Dilliard SA, Cheng Q, Siegwart DJ. On the mechanism of tissue-specific mRNA delivery by selective organ targeting nanoparticles. Proc Natl Acad Sci U S A. Dec 28 2021;118(52) doi:10.1073/pnas.2109256118

51. Cheng Q, Wei T, Farbiak L, Johnson LT, Dilliard SA, Siegwart DJ. Selective organ targeting (SORT) nanoparticles for tissue-specific mRNA delivery and CRISPR-Cas gene editing. Nat Nanotechnol. Apr 2020;15(4):313–320. doi:10.1038/s41565-020-0669-6

52. Hajj KA, Ball RL, Deluty SB, et al. Branched-Tail Lipid Nanoparticles Potently Deliver mRNA In Vivo due to Enhanced Ionization at Endosomal pH. Small. Feb 2019;15(6):e1805097. doi:10.1002/smll.201805097

53. Whitehead KA, Dorkin JR, Vegas AJ, et al. Degradable lipid nanoparticles with predictable in vivo siRNA delivery activity. Nat Commun. Jun 27 2014;5:4277. doi:10.1038/ncomms5277

54. Graham CH, Hawley TS, Hawley RG, et al. Establishment and Characterization of First Trimester Human Trophoblast Cells with Extended Lifespan. Experimental Cell Research. 1993;206:204–211.

55. Pattillo RA, Ruckert ACF, Hussa RO, Bernstein R, Delfs EM. The JAr cell line -- continuous human multihormone production and controls. Twenty-Second Annual Meeting of the Tissue Culture Association. 1971/04/01 1971;6(5):371–406. doi:10.1007/BF02619074

56. Hochberg A, Rachmilewitz J, Eldar-Geva T, Salant T, Schneider T, de Groot N. Differentiation of Choriocarcinoma Cell Line (JAr). Cancer research (Chicago, Ill). 1992;52(13):3713–3717.

57. Pattillo RA, Gey GO, Delfs EM, Mattingly RF. Human horomone production in vitro. Science. 1968;159:1467–1469. doi:10.1126/science.159.3822.1467

58. Rothbauer M, Patel N, Gondola H, Siwetz M, Huppertz B, Ertl P. A comparative study of five physiological key parameters between four different human trophoblast-derived cell lines. Sci Rep. Jul 19 2017;7(1):5892. doi:10.1038/s41598-017-06364-z

59. Caniggia I, Mostachfi H, Winter J, et al. Hypoxia-inducible factor-1 mediates the biological effects of oxygen on human trophoblast differentiation through TGFbeta(3). J Clin Invest. Mar 2000;105(5):577–87. doi:10.1172/JCI8316

60. Fujii T, Nagamatsu T, Morita K, et al. Enhanced HIF2alpha expression during human trophoblast differentiation into syncytiotrophoblast suppresses transcription of placental growth factor. Sci Rep. Sep 29 2017;7(1):12455. doi:10.1038/s41598-017-12685-w

61. Saffer C, Olson G, Boggess KA, et al. Determination of placental growth factor (PlGF) levels in healthy pregnant women without signs or symptoms of preeclampsia. Pregnancy Hypertens. Apr 2013;3(2):124–32. doi:10.1016/j.preghy.2013.01.004

62. Wang Y, Zhao S. Vascular Biology of the Placenta. Cell Types of the Placenta. Morgan & Claypool Life Sciences; 2010:chap 4.

63. Snegovskikh V, Hodgson E, Wehrum M, et al. 400: Human chorionic gonadotrophin (hCG) is produced by both syncytiotrophoblast and cytotrophoblast cells: A paradigm shift. American Journal of Obstetrics & Gynecology. 2007;197(6):S120. doi:10.1016/j.ajog.2007.10.418

64. Orendi K, Gauster M, Moser G, Meiri H, Huppertz B. The choriocarcinoma cell line BeWo: syncytial fusion and expression of syncytium-specific proteins. Reproduction. Nov 2010;140(5):759–66. doi:10.1530/REP-10-0221

65. Orendi K, Kivity V, Sammar M, et al. Placental and trophoblastic in vitro models to study preventive and therapeutic agents for preeclampsia. Placenta. Feb 2011;32 Suppl:S49–54. doi:10.1016/j.placenta.2010.11.023

66. Wang HL, Liang N, Huang DX, et al. The effects of high-density lipoprotein and oxidized high-density lipoprotein on forskolin-induced syncytialization of BeWo cells. Placenta. Jan 1 2021;103:199–205. doi:10.1016/j.placenta.2020.10.024

67. Abostait A, Tyrrell J, Abdelkarim M, et al. Placental Nanoparticle Uptake-On-a-Chip: The Impact of Trophoblast Syncytialization and Shear Stress. Mol Pharm. Sep 2 2022;doi:10.1021/acs.molpharmaceut.2c00216

68. Wice B, Menton D, Geuze H, Schwartz AL. Modulators of Cyclic AMP Metabolism Induce Syncytiotrophoblast Formation In Vitro. Experimental Cell Research. 1990;186:306–316.

69. Pidoux G, Gerbaud P, Gnidehou S, et al. ZO-1 is involved in trophoblastic cell differentiation in human placenta. Am J Physiol Cell Physiol. Jun 2010;298(6):C1517–26. doi:10.1152/ajpcell.00484.2008

70. Yang M, Chen Y, Chen L, et al. miR-15b-AGO2 play a critical role in HTR8/SVneo invasion and in a model of angiogenesis defects related to inflammation. Placenta. May 2016;41:62–73. doi:10.1016/j.placenta.2016.03.007

71. Ding GC, Chen M, Wang YX, et al. MicroRNA-128a-induced apoptosis in HTR-8/SVneo trophoblast cells contributes to pre-eclampsia. Biomed Pharmacother. Jul 2016;81:63–70. doi:10.1016/j.biopha.2016.03.040

72. Munaut C, Lorquet S, Pequeux C, et al. Hypoxia is responsible for soluble vascular endothelial growth factor receptor-1 (VEGFR-1) but not for soluble endoglin induction in villous trophoblast. Hum Reprod. Jun 2008;23(6):1407–15. doi:10.1093/humrep/den114

73. Cole LA. New discoveries on the biology and detection of human chorionic gonadotropin. Reprod Biol Endocrinol. Jan 26 2009;7:8. doi:10.1186/1477-7827-7-8

74. Suzuki H, Ohkuchi A, Matsubara S, et al. Effect of recombinant placental growth factor 2 on hypertension induced by full-length mouse soluble fms-like tyrosine kinase 1 adenoviral vector in pregnant mice. Hypertension. Nov 2009;54(5):1129–35. doi:10.1161/HYPERTENSIONAHA.109.134668

75. Spradley FT, Tan AY, Joo WS, et al. Placental Growth Factor Administration Abolishes Placental Ischemia-Induced Hypertension. Hypertension. Apr 2016;67(4):740–7. doi:10.1161/HYPERTENSIONAHA.115.06783

76. Yu J, Jia J, Guo X, Chen R, Feng L. Modulating circulating sFlt1 in an animal model of preeclampsia using PAMAM nanoparticles for siRNA delivery. Placenta. Oct 2017;58:1–8. doi:10.1016/j.placenta.2017.07.360

77. Makris A, Yeung KR, Lim SM, et al. Placental Growth Factor Reduces Blood Pressure in a Uteroplacental Ischemia Model of Preeclampsia in Nonhuman Primates. Hypertension. Jun 2016;67(6):1263–72. doi:10.1161/HYPERTENSIONAHA.116.07286

78. McCracken SA, Seeho SKM, Carrodus T, et al. Dysregulation of Oxygen Sensing/Response Pathways in Pregnancies Complicated by Idiopathic Intrauterine Growth Restriction and Early-Onset Preeclampsia. Int J Mol Sci. Mar 2 2022;23(5) doi:10.3390/ijms23052772

79. Rath G, Aggarwal R, Jawanjal P, Tripathi R, Batra A. HIF-1 Alpha and Placental Growth Factor in Pregnancies Complicated With Preeclampsia: A Qualitative and Quantitative Analysis. J Clin Lab Anal. Jan 2016;30(1):75–83. doi:10.1002/jcla.21819

80. Roland CS, Hu J, Ren CE, et al. Morphological changes of placental syncytium and their implications for the pathogenesis of preeclampsia. Cell Mol Life Sci. Jan 2016;73(2):365–76. doi:10.1007/s00018-015-2069-x

81. Alsat E, Wyplosz P, Malassiné A, et al. Hypoxia impairs cell fusion and differentiation process in human cytotrophoblast, in vitro. Journal of Cellular Physiology. 1996;168(2):346–353. doi:10.1002/(sici)1097-4652(199608)168:2<346::Aid-jcp13>3.0.Co;2-1

82. Kudo Y, Boyd CA, Sargent IL, Redman CW. Hypoxia alters expression and function of syncytin and its receptor during trophoblast cell fusion of human placental BeWo cells: implications for impaired trophoblast syncytialisation in pre-eclampsia. Biochim Biophys Acta. May 20 2003;1638(1):63–71. doi:10.1016/s0925-4439(03)00043-7

83. Jaremek A, Shaha S, Jeyarajah MJ, et al. Genome-Wide Analysis of Hypoxia-Inducible Factor Binding Reveals Targets Implicated in Impaired Human Placental Syncytiotrophoblast Formation under Low Oxygen. Am J Pathol. Jul 2023;193(7):846–865. doi:10.1016/j.ajpath.2023.03.006

84. Colson A, Depoix CL, Baldin P, Hubinont C, Sonveaux P, Debieve F. Hypoxia-inducible factor 2 alpha impairs human cytotrophoblast syncytialization: New insights into placental dysfunction and fetal growth restriction. FASEB J. Nov 2020;34(11):15222–15235. doi:10.1096/fj.202001681R

85. Albers RE, Kaufman MR, Natale BV, et al. Trophoblast-Specific Expression of Hif-1alpha Results in Preeclampsia-Like Symptoms and Fetal Growth Restriction. Sci Rep. Feb 26 2019;9(1):2742. doi:10.1038/s41598-019-39426-5

86. Delehedde C, Even L, Midoux P, Pichon C, Perche F. Intracellular Routing and Recognition of Lipid-Based mRNA Nanoparticles. Pharmaceutics. Jun 24 2021;13(7) doi:10.3390/pharmaceutics13070945

87. Li G, Wang Y, Cao G, et al. Hypoxic stress disrupts HGF/Met signaling in human trophoblasts: implications for the pathogenesis of preeclampsia. J Biomed Sci. Feb 3 2022;29(1):8. doi:10.1186/s12929-022-00791-5

88. Horst B, Pradhan S, Chaudhary R, et al. Hypoxia-induced inhibin promotes tumor growth and vascular permeability in ovarian cancers. Commun Biol. Jun 2 2022;5(1):536. doi:10.1038/s42003-022-03495-6

89. Liu F, Soares MJ, Audus KL. Permeability properties of monolayers of the human trophoblast cell line BeWo. American Journal of Physiology. 1997;273(5):C1596–C1604. doi:10.1152/ajpcell.1997.273.5.C1596

90. Kouthouridis S, Sotra A, Khan Z, Alvarado J, Raha S, Zhang B. Modeling the Progression of Placental Transport from Early-to Late-Stage Pregnancy by Tuning Trophoblast Differentiation and Vascularization. Adv Healthc Mater. Dec 2023;12(32):e2301428. doi:10.1002/adhm.202301428

91. Autiero M, Waltenberger J, Communi D, et al. Role of PlGF in the intra- and intermolecular cross talk between the VEGF receptors Flt1 and Flk1. Nature Medicine. 2003;9(7):936–943. doi:10.1038/nm884

92. Andrikos A, Andrikos D, Schmidt B, et al. Course of the sFlt-1/PlGF ratio in fetal growth restriction and correlation with biometric measurements, feto-maternal Doppler parameters and time to delivery. Arch Gynecol Obstet. Mar 2022;305(3):597–605. doi:10.1007/s00404-021-06186-5

93. Agrawal S, Shinar S, Cerdeira AS, Redman C, Vatish M. Predictive Performance of PlGF (Placental Growth Factor) for Screening Preeclampsia in Asymptomatic Women: A Systematic Review and Meta-Analysis. Hypertension. Nov 2019;74(5):1124–1135. doi:10.1161/HYPERTENSIONAHA.119.13360

94. Knudsen UB, Kronborg CS, von Dadelszen P, et al. A single rapid point-of-care placental growth factor determination as an aid in the diagnosis of preeclampsia. Pregnancy Hypertens. Jan 2012;2(1):8–15. doi:10.1016/j.preghy.2011.08.117

95. Jaremek A, Jeyarajah MJ, Jaju Bhattad G, Renaud SJ. Omics Approaches to Study Formation and Function of Human Placental Syncytiotrophoblast. Front Cell Dev Biol. 2021;9:674162. doi:10.3389/fcell.2021.674162

96. Heyes J, Palmer L, Bremner K, MacLachlan I. Cationic lipid saturation influences intracellular delivery of encapsulated nucleic acids. J Control Release. Oct 3 2005;107(2):276–87. doi:10.1016/j.jconrel.2005.06.014

