## Supplemental File for "Investigating the Impact of Hypoxia and Syncytialization on Lipid Nanoparticle-Mediated mRNA Delivery to Placental Cells"

*Postal: 201 Mullica Hill Rd, Glassboro, NJ, 08028

**Table of Contents**

**Figure S1.** Cell metabolic activity following LNP delivery.

**Figure S2.** Hypoxic cell culture set-up.

**Figure S3.** Cell metabolic activity following LNP delivery in normoxia and hypoxia.

**Table S1.** LNP Library Parameters and Characterization

**Table S2.** Main Effects Estimates from Definitive Screening Fit for Luc-LNP Delivery in HTR8 Cells

**Table S3.** Main Effects Estimates from Definitive Screening Fit for Luc-LNP Delivery in JAR Cells

**Table S4.** Ordinary Two-Way ANOVA Table for Grouped Lipid Analysis of Luc-LNP Delivery in HTR8 Cells

**Table S5.** Ordinary Two-Way ANOVA Table for Grouped Lipid Analysis of Luc-LNP Delivery in JAR Cells

**Table S6.** Ordinary Two-Way ANOVA Table for Grouped Lipid Analysis of Luc-LNP Delivery in BeWo Cells

**Table S7.** Additional Comparisons from Tukey Multiple Comparisons Test of HIF-1α Data in Hypoxic Cells

**Table S8.** Ordinary Two-Way ANOVA Table for HIF-1α Data

**Table S9.** Additional Comparisons from Tukey Multiple Comparisons Test of PlGF Data in Normoxic and Hypoxic Cells

**Table S10.** Ordinary Two-Way ANOVA Table for PlGF Data

**Table S11.** Ordinary Two-Way ANOVA Table for MFI in HTR8 Cells 2 Hours after GFP-LNP Delivery

**Table S12.** Ordinary Two-Way ANOVA Table for MFI in HTR8 Cells 24 Hours after GFP-LNP Delivery

**Table S13.** Ordinary Two-Way ANOVA Table for MFI in JAR Cells 2 Hours after GFP-LNP Delivery

**Table S14.** Ordinary Two-Way ANOVA Table for MFI in JAR Cells 24 Hours after GFP-LNP Delivery

**Table S15.** Ordinary Two-Way ANOVA Table for MFI in BeWo Cells 2 Hours after GFP-LNP Delivery

**Table S16.** Ordinary Two-Way ANOVA Table for MFI in BeWo Cells 24 Hours after GFP-LNP Delivery

**Table S17.** Ordinary Two-Way ANOVA Table for hCG Secretion from JAR Cells

**Table S18.** Ordinary Two-Way ANOVA Table for PlGF Secretion from JAR Cells

**Table S19.** Ordinary Two-Way ANOVA Table for hCG Secretion from BeWo Cells

**Table S20.** Ordinary Two-Way ANOVA Table for PlGF Secretion from BeWo Cells

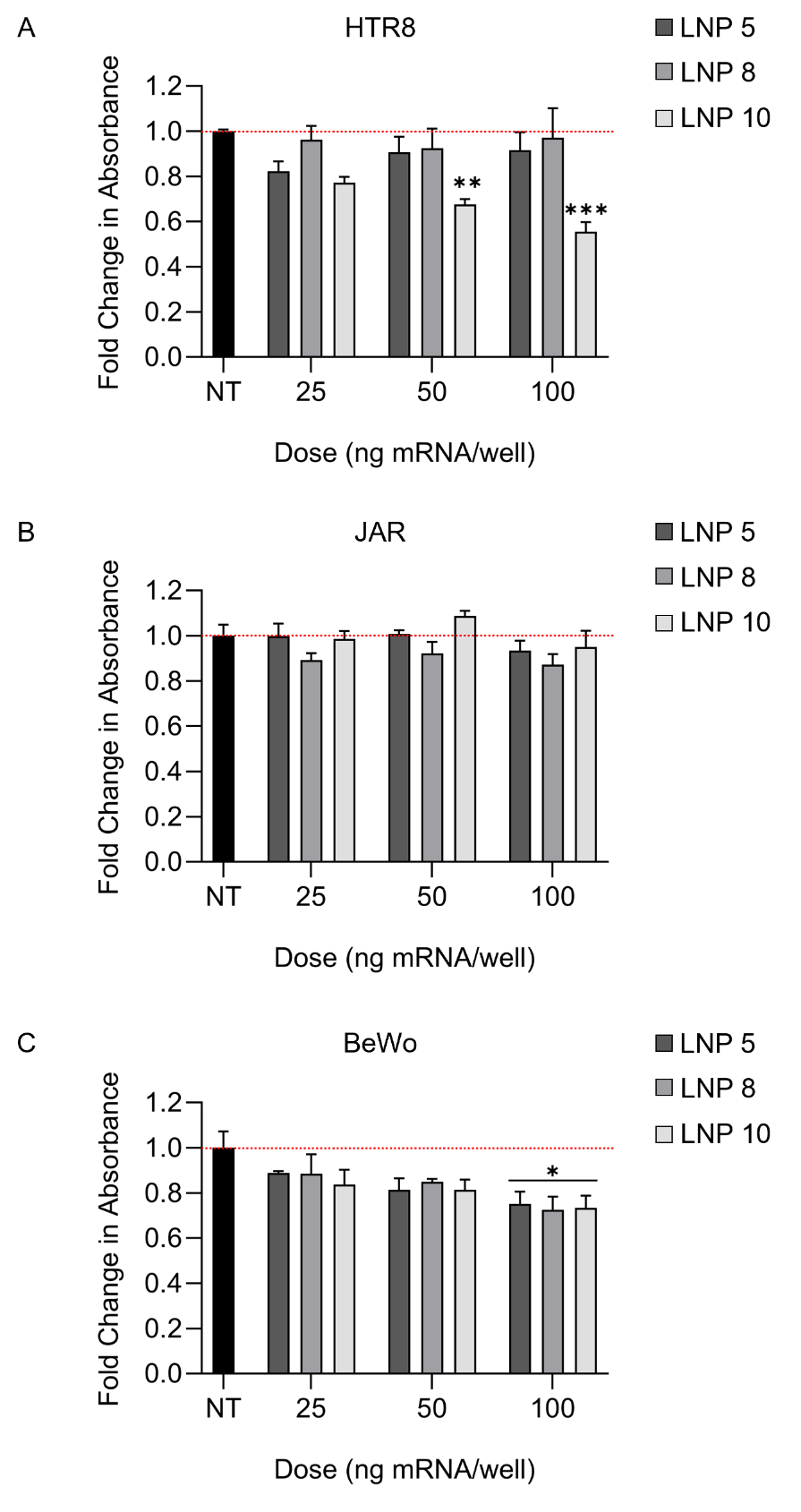

**Figure S1.** Cell metabolic activity following LNP delivery. (A) HTR8, (B) JAR, and (C) BeWo metabolic activity following delivery with select Luc-LNPs from library (LNP 5, LNP 8, LNP 10) at varying doses of mRNA. *p<0.05, **p<0.01,***p<0.001 by two-way ANOVA with posthoc Dunnett’s comparing groups to the non-treated “NT” group within each cell line.

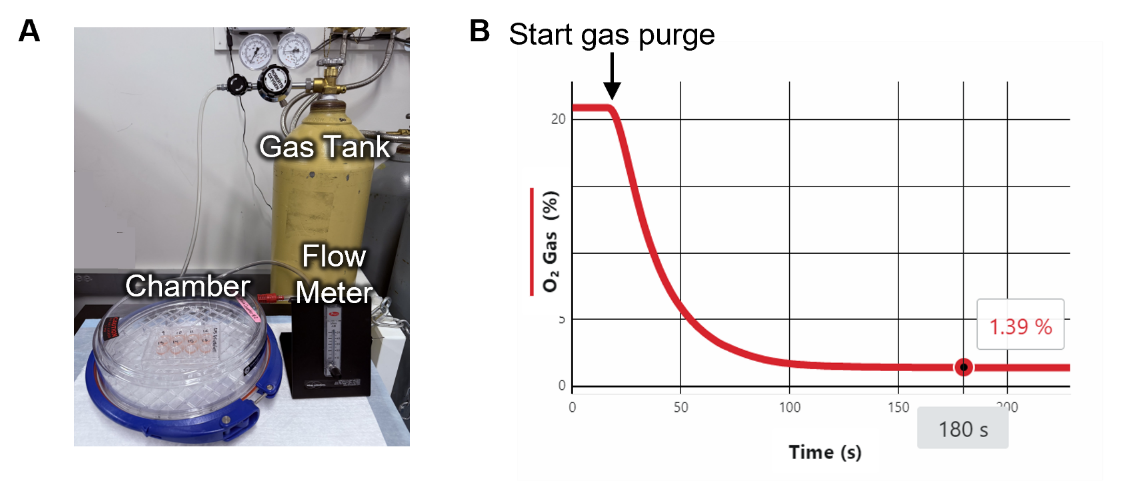

**Figure S2.** Hypoxic cell culture set-up. (A) Cell culture chamber used to create a low oxygen environment. (B) Oxygen concentration in the chamber versus time.

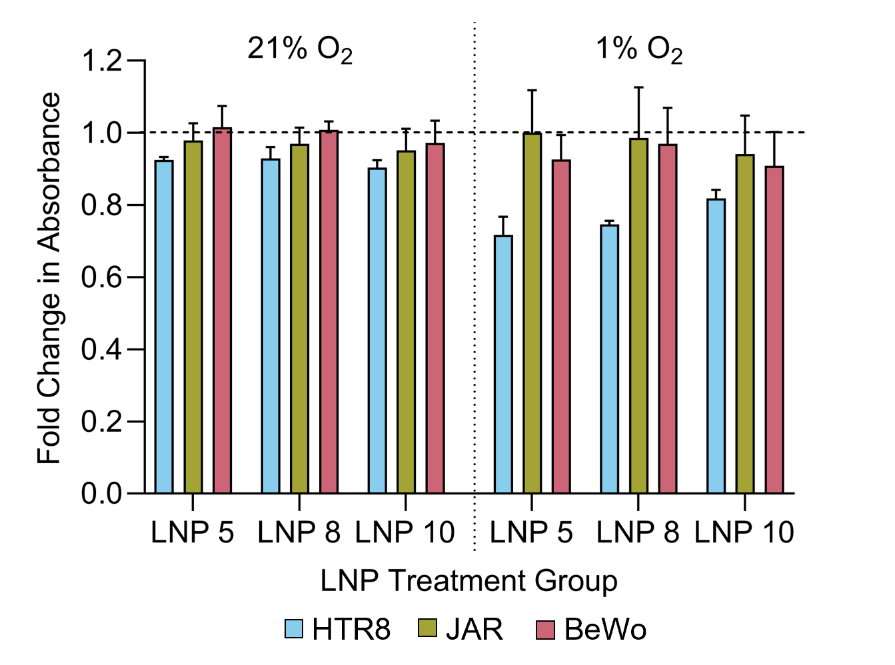

**Figure S3.** Cell metabolic activity in normoxia and hypoxia following delivery with select LNPs from the library (LNP 5, LNP 8, LNP 10) at 25 ng mRNA/well.

**Table S1.** LNP Library Parameters and Characterization

| **LNP** | **Ion. ID** | **Phos.ID** | **Ion. (%)** | **Phos. (%)** | | **PEG (%)** | **Chol. (%)** | **HDD (nm)** | **PDI** | **EE (%)** | **pKa** |
| --- | --- | --- | --- | --- | --- | --- | --- | --- | --- | --- | --- |
| A1 | C12 | DSPC | 45 | 22 | 3.5 | | 29.5 | 114.8 | 0.141 | 60.5 | 6.098 |
| A2 | C12 | DSPC | 25 | 10 | 3.5 | | 61.5 | 90.3 | 0.118 | 66.6 | 6.358 |
| A3 | C12 | DOPE | 45 | 10 | 3.5 | | 41.5 | 104.9 | 0.0541 | 41.5 | 5.878 |
| A4 | MC3 | DOPE | 45 | 22 | 1.5 | | 31.5 | 114.8 | 0.0734 | 42.0 | 6.806 |
| A5 | C12 | DOPE | 25 | 22 | 2.5 | | 50.5 | 141.4 | 0.0622 | 55.0 | 5.795 |
| A6 | MC3 | DOPE | 25 | 10 | 1.5 | | 63.5 | 126.4 | 0.118 | 87.1 | 6.96 |
| A7 | MC3 | DSPC | 45 | 10 | 1.5 | | 43.5 | 143.7 | 0.183 | 81.3 | 6.382 |
| A8 | C12 | DOPE | 25 | 22 | 3.5 | | 49.5 | 125.4 | 0.0837 | 73.7 | 5.798 |
| A9 | MC3 | DOPE | 45 | 16 | 3.5 | | 35.5 | 110.3 | 0.0969 | 62.2 | 6.646 |
| A10 | C12 | DOPE | 35 | 10 | 1.5 | | 53.5 | 153.9 | 0.0269 | 63.7 | 6.308 |
| A11 | MC3 | DOPE | 25 | 10 | 3.5 | | 61.5 | 109.7 | 0.0873 | 82.0 | 7.111 |
| A12 | MC3 | DSPC | 35 | 16 | 2.5 | | 46.5 | 96.82 | 0.170 | 74.5 | 6.42 |
| A13 | MC3 | DSPC | 25 | 22 | 1.5 | | 51.5 | 111.8 | 0.129 | 81.1 | 6.64 |
| A14 | C12 | DOPE | 35 | 16 | 2.5 | | 46.5 | 115 | 0.100 | 58.5 | 5.448 |
| A15 | C12 | DSPC | 45 | 22 | 1.5 | | 31.5 | 121.6 | 0.121 | 73.7 | 5.298 |
| A16 | C12 | DSPC | 25 | 16 | 1.5 | | 57.5 | 103.4 | 0.134 | 80.6 | 6.533 |
| A17 | MC3 | DSPC | 35 | 22 | 3.5 | | 39.5 | 107.4 | 0.163 | 74.1 | 6.68 |
| A18 | MC3 | DSPC | 45 | 10 | 2.5 | | 42.5 | 111.3 | 0.147 | 78.1 | 6.407 |

Ionizable lipid type, Ion.ID; C12-200, C12; DLin-MC3-DMA, MC3; Phospholipid type, Phos.ID; 1,2-dioleoyl-sn-glycero-3-phosphoethanolamine, DOPE; 1,2 distearoyl-sn-glycero-3-phosphocholine, DSPC; Ionizable lipid, Ion.; Phospholipid, Phos.; Cholesterol, Chol.; Poly(ethylene) glycol, PEG; Hydrodynamic Diameter, HDD; Polydispersity index, PDI; Encapsulation Efficiency, EE; Apparent pKa, pKa; Zeta Potential, Zeta;

**Table S2.** Main Effects Estimates from Definitive Screening Fit for Luc-LNP Delivery in HTR8 Cells

| Term | Estimate | Std Error | t Ratio | Prob >\|t\| | Statistic | Value |
| --- | --- | --- | --- | --- | --- | --- |
| Phospholipid Type | 9880.1 | 1887.4 | 5.2346 | 0.0012 | **RMSE** | 7944 |
| Phospholipid Amount | 6726.4 | 2140.2 | 3.1429 | 0.0163 | **DF** | 7 |

**Table S3.** Main Effects Estimates from Definitive Screening Fit for Luc-LNP Delivery in JAR Cells

| Term | Estimate | Std Error | t Ratio | Prob >\|t\| | Statistic | Value |
| --- | --- | --- | --- | --- | --- | --- |
| Phospholipid Type | 8322.2 | 2594 | 3.2082 | 0.0125 | **RMSE** | 11006 |
|  |  |  |  |  | **DF** | 8 |

**Table S4.** Ordinary Two-Way ANOVA Table for Grouped Lipid Analysis of Luc-LNP Delivery in HTR8 Cells

| **ANOVA table** | **F (DFn, DFd)** | **DF** | **% of total variation** | **P value** | **P Value Summary** |
| --- | --- | --- | --- | --- | --- |
| **Interaction** | F (1, 14) = 0.9442 | 1 | 3.526 | 0.3477 | ns |
| **Ionizable Lipid** | F (1, 14) = 0.8883 | 1 | 3.317 | 0.3619 | ns |
| **Phospholipid** | F (1, 14) = 10.13 | 1 | 37.84 | 0.0066 | ** |

**Table S5.** Ordinary Two-Way ANOVA Table for Grouped Lipid Analysis of Luc-LNP Delivery in JAR Cells

| **ANOVA table** | **F (DFn, DFd)** | **DF** | **% of total variation** | **P value** | **P Value Summary** |
| --- | --- | --- | --- | --- | --- |
| **Interaction** | F (1, 14) = 1.773 | 1 | 6.355 | 0.2043 | ns |
| **Ionizable Lipid** | F (1, 14) = 0.9665 | 1 | 3.464 | 0.3422 | ns |
| **Phospholipid** | F (1, 14) = 10.31 | 1 | 36.95 | 0.0063 | ** |

**Table S6.** Ordinary Two-Way ANOVA Table for Grouped Lipid Analysis of Luc-LNP Delivery in BeWo Cells

| **ANOVA table** | **F (DFn, DFd)** | **DF** | **% of total variation** | **P value** | **P Value Summary** |
| --- | --- | --- | --- | --- | --- |
| **Interaction** | F (1, 14) = 3.335 | 1 | 16.33 | 0.0892 | ns |
| **Ionizable Lipid** | F (1, 14) = 2.062 | 1 | 10.09 | 0.1730 | ns |
| **Phospholipid** | F (1, 14) = 0.7207 | 1 | 3.528 | 0.4102 | ns |

**Table S7.** Additional Comparisons from Tukey Multiple Comparisons Test of HIF-1α Data in Hypoxic Cells

| **Group** | **P value** | **P Value Summary** |
| --- | --- | --- |
| **JAR 24 vs. JAR 48** | <0.0001 | **** |
| **JAR 24 vs. JAR 72** | <0.0001 | **** |
| **JAR 48 vs. JAR 72** | 0.2312 | ns |
| **BeWo 24 vs. BeWo 48** | <0.0001 | **** |
| **BeWo 24 vs. BeWo 72** | <0.0001 | **** |
| **BeWo 48 vs. BeWo 72** | 0.2312 | ns |

**Table S8.** Ordinary Two-Way ANOVA Table for HIF-1α Data

| **ANOVA table** | **F (DFn, DFd)** | **DF** | **% of total variation** | **P value** | **P Value Summary** |
| --- | --- | --- | --- | --- | --- |
| **Time** | F (2, 85) = 28.03 | 2 | 18.53 | <0.0001 | **** |
| **Cell and Oxygen** | F (4, 85) = 51.91 | 4 | 68.63 | <0.0001 | **** |

**Table S9.** Additional Comparisons from Tukey Multiple Comparisons Test of PlGF Data in Normoxic and Hypoxic Cells

| **Group** | **P value** | **P Value Summary** |
| --- | --- | --- |
| **21%: JAR 24 vs. JAR 48** | <0.0001 | **** |
| **21%: JAR 24 vs. JAR 72** | <0.0001 | **** |
| **21%: JAR 48 vs. JAR 72** | <0.0001 | **** |
| **21%: BeWo 24 vs. BeWo 48** | 0.0006 | *** |
| **21%: BeWo 24 vs. BeWo 72** | <0.0001 | **** |
| **21%: BeWo 48 vs. BeWo 72** | 0.2312 | ns |
| **1%: JAR 24 vs. JAR 48** | 0.7324 | ns |
| **1%: JAR 24 vs. JAR 72** | <0.0001 | **** |
| **1%: JAR 48 vs. JAR 72** | <0.0001 | **** |
| **1%: BeWo 24 vs. BeWo 48** | 0.5178 | ns |
| **1%: BeWo 24 vs. BeWo 72** | 0.0009 | *** |
| **1%: BeWo 48 vs. BeWo 72** | 0.5817 | ns |

**Table S10.** Ordinary Two-Way ANOVA Table for PlGF Data

| **ANOVA table** | **F (DFn, DFd)** | **DF** | **% of total variation** | **P value** | **P Value Summary** |
| --- | --- | --- | --- | --- | --- |
| **Interaction** | F (10, 54) = 21.38 | 10 | 16.93 | <0.0001 | **** |
| **Time** | F (2, 54) = 165.1 | 2 | 26.15 | <0.0001 | **** |
| **Cell and Oxygen** | F (5, 54) = 132.9 | 5 | 52.64 | <0.0001 | **** |

**Table S11.** Ordinary Two-Way ANOVA Table for MFI in HTR8 Cells 2 Hours after GFP-LNP Delivery

| **ANOVA table** | **F (DFn, DFd)** | **DF** | **% of total variation** | **P value** | **P Value Summary** |
| --- | --- | --- | --- | --- | --- |
| **Interaction** | F (2, 12) = 0.03396 | 2 | 0.2437 | 0.9667 | ns |
| **GFP-LNP** | F (2, 12) = 7.771 | 2 | 55.76 | 0.0068 | ** |
| **Oxygen** | F (1, 12) = 0.2646 | 1 | 0.9491 | 0.6163 | ns |

**Table S12.** Ordinary Two-Way ANOVA Table for MFI in HTR8 Cells 24 Hours after GFP-LNP Delivery

| **ANOVA table** | **F (DFn, DFd)** | **DF** | **% of total variation** | **P value** | **P Value Summary** |
| --- | --- | --- | --- | --- | --- |
| **Interaction** | F (2, 12) = 0.05570 | 2 | 0.5485 | 0.9461 | ns |
| **GFP-LNP** | F (2, 12) = 3.782 | 2 | 37.24 | 0.0532 | ns |
| **Oxygen** | F (1, 12) = 0.6365 | 1 | 3.133 | 0.4405 | ns |

**Table S13.** Ordinary Two-Way ANOVA Table for MFI in JAR Cells 2 Hours after GFP-LNP Delivery

| **ANOVA table** | **F (DFn, DFd)** | **DF** | **% of total variation** | **P value** | **P Value Summary** |
| --- | --- | --- | --- | --- | --- |
| **Interaction** | F (2, 12) = 0.7141 | 2 | 6.995 | 0.5093 | ns |
| **GFP-LNP** | F (2, 12) = 0.5322 | 2 | 5.213 | 0.6006 | ns |
| **Oxygen** | F (1, 12) = 5.925 | 1 | 29.02 | 0.0315 | * |

**Table S14.** Ordinary Two-Way ANOVA Table for MFI in JAR Cells 24 Hours after GFP-LNP Delivery

| **ANOVA table** | **F (DFn, DFd)** | **DF** | **% of total variation** | **P value** | **P Value Summary** |
| --- | --- | --- | --- | --- | --- |
| **Interaction** | F (2, 12) = 0.4023 | 2 | 3.213 | 0.6775 | ns |
| **GFP-LNP** | F (2, 12) = 5.703 | 2 | 45.55 | 0.0182 | * |
| **Oxygen** | F (1, 12) = 0.8311 | 1 | 3.319 | 0.3799 | ns |

**Table S15.** Ordinary Two-Way ANOVA Table for MFI in BeWo Cells 2 Hours after GFP-LNP Delivery

| **ANOVA table** | **F (DFn, DFd)** | **DF** | **% of total variation** | **P value** | **P Value Summary** |
| --- | --- | --- | --- | --- | --- |
| **Interaction** | F (2, 11) = 0.1615 | 2 | 1.667 | 0.8529 | ns |
| **GFP-LNP** | F (2, 11) = 0.9767 | 2 | 10.09 | 0.4069 | ns |
| **Oxygen** | F (1, 11) = 5.745 | 1 | 29.66 | 0.0354 | * |

**Table S16.** Ordinary Two-Way ANOVA Table for MFI in BeWo Cells 24 Hours after GFP-LNP Delivery

| **ANOVA table** | **F (DFn, DFd)** | **DF** | **% of total variation** | **P value** | **P Value Summary** |
| --- | --- | --- | --- | --- | --- |
| **Interaction** | F (2, 11) = 7.064 | 2 | 6.238 | 0.0106 | * |
| **GFP-LNP** | F (2, 11) = 66.21 | 2 | 58.47 | <0.0001 | **** |
| **Oxygen** | F (1, 11) = 86.45 | 1 | 38.17 | <0.0001 | **** |

**Table S17.** Ordinary Two-Way ANOVA Table for hCG Secretion from JAR Cells

| **ANOVA table** | **F (DFn, DFd)** | **DF** | **% of total variation** | **P value** | **P Value Summary** |
| --- | --- | --- | --- | --- | --- |
| **Interaction** | F (1, 20) = 7.916 | 1 | 3.112 | 0.0107 | * |
| **Forskolin Treatment** | F (1, 20) = 193.7 | 1 | 76.13 | <0.0001 | **** |
| **Oxygen** | F (1, 20) = 32.81 | 1 | 12.89 | <0.0001 | **** |

**Table S18.** Ordinary Two-Way ANOVA Table for PlGF Secretion from JAR Cells

| **ANOVA table** | **F (DFn, DFd)** | **DF** | **% of total variation** | **P value** | **P Value Summary** |
| --- | --- | --- | --- | --- | --- |
| **Interaction** | F (1, 20) = 29.42 | 1 | 5.934 | <0.0001 | **** |
| **Forskolin Treatment** | F (1, 20) = 383.9 | 1 | 77.43 | <0.0001 | **** |
| **Oxygen** | F (1, 20) = 62.48 | 1 | 12.60 | <0.0001 | **** |

**Table S19.** Ordinary Two-Way ANOVA Table for hCG Secretion from BeWo Cells

| **ANOVA table** | **F (DFn, DFd)** | **DF** | **% of total variation** | **P value** | **P Value Summary** |
| --- | --- | --- | --- | --- | --- |
| **Interaction** | F (1, 20) = 16.63 | 1 | 11.72 | 0.0006 | *** |
| **Forskolin Treatment** | F (1, 20) = 84.04 | 1 | 59.22 | <0.0001 | **** |
| **Oxygen** | F (1, 20) = 21.24 | 1 | 14.97 | 0.0002 | *** |

**Table S20.** Ordinary Two-Way ANOVA Table for PlGF Secretion from BeWo Cells

| **ANOVA table** | **F (DFn, DFd)** | **DF** | **% of total variation** | **P value** | **P Value Summary** |
| --- | --- | --- | --- | --- | --- |
| **Interaction** | F (1, 20) = 91.44 | 1 | 7.995 | <0.0001 | **** |
| **Forskolin Treatment** | F (1, 20) = 857.7 | 1 | 74.99 | <0.0001 | **** |
| **Oxygen** | F (1, 20) = 174.5 | 1 | 15.26 | <0.0001 | **** |

**Table S21.** Ordinary Two-Way ANOVA Table for MFI in BeWo Cells 24 Hours after GFP-LNP Delivery

| **ANOVA table** | **F (DFn, DFd)** | **DF** | **% of total variation** | **P value** | **P Value Summary** |
| --- | --- | --- | --- | --- | --- |
| **Interaction** | F (1, 12) = 1.332 | 1 | 4.622 | 0.2709 | ns |
| **Forskolin Treatment** | F (1, 12) = 4.548 | 1 | 15.78 | 0.0543 | ** |
| **Oxygen** | F (1, 12) = 10.94 | 1 | 37.96 | 0.0062 | ns |

**Table S22.** Ordinary Three-Way ANOVA Table for PlGF Secretion from BeWo Cells 24 Hours after PlGF-LNP Delivery

| **ANOVA table** | **F (DFn, DFd)** | **DF** | **% of total variation** | **P value** | | **P Value Summary** |
| --- | --- | --- | --- | --- | --- | --- |
| **LNPs** | F (1, 61) = 283.7 | 1 | 63.81 | <0.0001 | **** | |
| **Oxygen** | F (1, 61) = 15.09 | 1 | 3.394 | 0.0003 | *** | |
| **Forskolin Treatment** | F (1, 61) = 40.25 | 1 | 9.051 | <0.0001 | **** | |
| **LNPs x Oxygen** | F (1, 61) = 1.831 | 1 | 0.4117 | 0.1810 | ns | |
| **LNPs x Forskolin Treatment** | F (1, 61) = 15.80 | 1 | 3.554 | 0.0002 | *** | |
| **Oxygen x Forskolin Treatment** | F (1, 61) = 4.926 | 1 | 1.108 | 0.0302 | * | |
| **LNPs x Oxygen x Forskolin Treatment** | F (1, 61) = 0.003586 | 1 | 0.0008064 | 0.9524 | ns | |

**Table S23.** Ordinary Two-Way ANOVA Table for fold change in PlGF Secretion from BeWo Cells 24 Hours after PlGF-LNP Delivery

| **ANOVA table** | **F (DFn, DFd)** | **DF** | **% of total variation** | **P value** | **P Value Summary** |
| --- | --- | --- | --- | --- | --- |
| **Interaction** | F (1, 29) = 3.175 | 1 | 2.245 | 0.0852 | ns |
| **Forskolin Treatment** | F (1, 29) = 14.11 | 1 | 9.978 | 0.0008 | *** |
| **Oxygen** | F (1, 29) = 95.15 | 1 | 67.28 | <0.0001 | **** |
